# Inferring cell cycle phases from a partially temporal network of protein interactions

**DOI:** 10.1101/2021.03.26.437187

**Authors:** Maxime Lucas, Arthur Morris, Alex Townsend-Teague, Laurent Tichit, Bianca H. Habermann, Alain Barrat

**Author notes:** Correspondence during submission should be addressed to: Bianca H. Habermann Aix-Marseille University, CNRS, IBDM UMR7288 Turing Center for Living Systems (CENTURI) Parc Scientifique de Luminy, Case 907, 163, Avenue de Luminy, 13009 Marseille, France. equal contribution (Laurent Tichit, Bianca H. Habermann, Alain Barrat).

## Abstract

The temporal organisation of biological systems into phases and subphases is often crucial to their functioning. Identifying this multiscale organisation can yield insight into the underlying biological mechanisms at play. To date, however, this identification requires *a priori* biological knowledge of the system under study. Here, we recover the temporal organisation of the cell cycle of budding yeast into phases and subphases, in an automated way. To do so, we model the cell cycle as a partially temporal network of protein-protein interactions (PPIs) by combining a traditional static PPI network with protein concentration or RNA expression time series data. Then, we cluster the snapshots of this temporal network to infer phases, which show good agreement with our biological knowledge of the cell cycle. We systematically test the robustness of the approach and investigate the effect of having only partial temporal information. Our results show for the first time that a temporal network with only partial temporal information, i.e. for some of the PPIs, is sufficient to infer the temporal organization of a system. The generality of the method makes it suitable for application to other, less well-known biological systems for which the temporal organisation of processes plays an important role.

## Introduction

Many biological systems go through successive phases, or states, over multiple time and space scales. These phases, and their order in time, are often crucial to the functioning of these systems and even determine their fate. To understand or predict these phases would yield crucial insight into the temporal organisation of such systems.

A particularly important example of such a process is the cell cycle, at the heart of all biological development: the precise timing of molecular events is crucial for the proper execution of cell division. Indeed, the cell typically progresses through 4 macroscopic phases, before eventually dividing into two cells (Figure 1). Historically, these cell cycle phases were first determined by A. Howard and S.P. Pelc at the cellular scale by analysing the proliferation of bean root cells (Howard and Pelc 1953). The cycle starts with a first gap phase (G1), in which the cell grows and needs to reach a certain size to enter the next phase, where DNA is synthesised (S), followed by a second gap phase (G2), and finally mitosis (M), in which the duplicated chromosome set is divided and equally distributed into two daughter cells. These 4 phases can then be further divided into subphases, or physiological processes. For example, mitosis is composed of prophase, metaphase, anaphase, and telophase.

**Figure 1:**
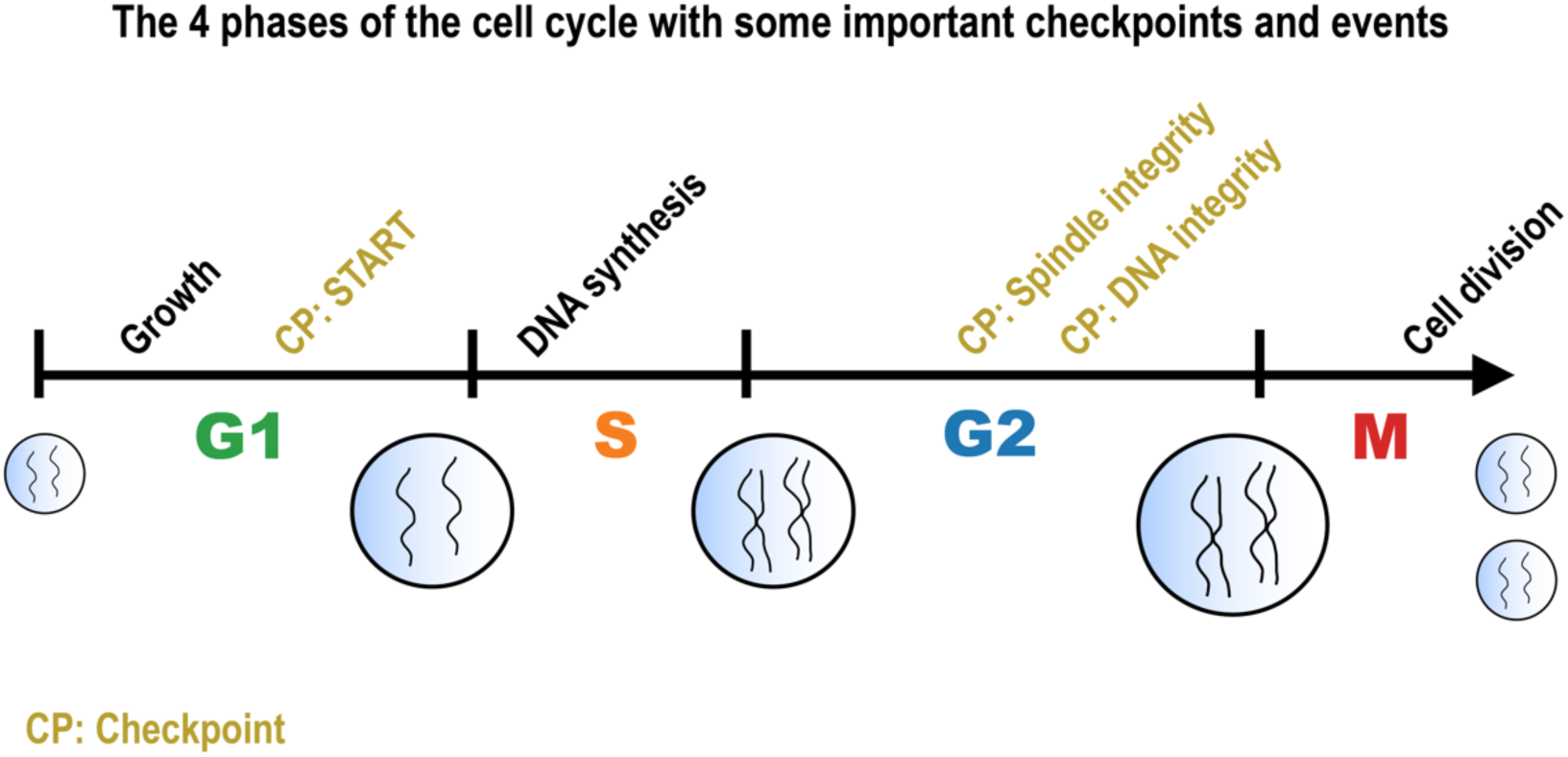
The cell cycle consists of a succession of physiological processes, mediated by protein-protein interactions leading to the division of the cell. The ordering of these interactions is crucial, and is ensured by molecular checkpoints (yellow). The cycle is divided into 4 main phases: Gap1-phase (G1), Synthesis-phase (S), Gap2-phase (G2), and Mitosis (M).

In the past 70 years, extensive cell cycle research identified the molecular events driving the behaviour of the cell in each of these 4 phases, see reviews by (Nasmyth 1993, 1996; Koch and Nasmyth 1994; Murray 2004; Vodermaier 2004). In particular, the transcriptionally controlled cyclins and the cyclin-dependent kinases were found to regulate a multitude of cell cycle events. A cellular destruction machinery was found to complement these proteins by degrading cell cycle regulators at specific times in the cell cycle. A set of core cell cycle proteins thus regulate the cell cycle and its phases by interacting with each other in a precisely timed manner. These interactions can be of different types: stable and long-term, transient and leading, e.g., to protein phosphorylation, de-phosphorylation, ubiquitination and/or degradation. They drive the progression of the cell cycle. As so-called molecular checkpoints, some ensure that one phase is complete before the cell enters the next phase. Not fulfilling a checkpoint can stop the progression of the cell cycle and change the fate of the cell.

The complexity of these interactions between proteins is often encoded in a network representation (Newman 2018). In such biological networks, nodes represent proteins and edges represent protein-protein interactions (PPIs) (Alon 2003; Pavlopoulos et al. 2011). The cell cycle has for instance been represented as a network of protein-protein interactions in the KEGG pathway resource (Kanehisa and Goto 2000). These network representations are however static; they do not include any information about the time-varying interactions between cell cycle regulators.

Nonetheless, over the past few years, temporal information about biological systems has become more widely available in the form of time series recordings thanks to high-throughput techniques such as RNA-sequencing and high-throughput proteomics. This has opened the door to the investigation of the temporal dynamics of various systems. In addition, mathematical models of specific biological systems have been proposed, based on a priori knowledge. These models typically describe the dynamics of gene expression levels (Chen et al. 1998) or protein concentrations (Chen et al. 2004), either as continuously evolving variables or using binary values (Traynard et al. 2016).

To incorporate such temporal information into the representation of protein-protein interactions, a natural framework is that of temporal networks (Holme and Saramäki 2012; Holme 2015). In a temporal network of PPIs, nodes represent proteins, and interactions between them are represented by time-varying edges. Temporal network theory has been used successfully in areas ranging from social interactions (Miritello et al. 2011; Saramäki and Moro 2015; Gelardi et al. 2020) to neuroscience (Pedreschi et al. 2020; Lopes et al. 2021). However, despite calls to use it more in biology (Przytycka et al. 2010), it is still underused to study, e.g., PPI networks. Existing studies mostly integrate gene expression data from microarray or RNA-sequencing to a static PPI network (Li et al. 2018) to identify active subnetworks (Pierrelée et al. 2020) and functional modules (Komurov and White 2007; Chechik et al. 2008; Li et al. 2012; Ou-Yang et al. 2014).

Here, we go further and fully take advantage of the temporal network framework and develop an automated pipeline to investigate the temporal organisation of the cell cycle across a range of time scales, starting from time series data of protein-protein interactions. To this end, we build a representation of the cell cycle as a temporal network of protein-protein interactions by incorporating time series data, obtained from a mathematical model of the cell cycle, into a static network of PPIs. We then build on a recent method introduced in (Masuda and Holme 2019) to infer phases by clustering snapshots of the temporal network. We finally validate these phases against ground truth biological knowledge. Within this pipeline, we present in particular three main methodological advances: (1) we analyse the phase inference results at multiple temporal scales, uncovering several additional subphases in the cell cycle, (2) we analyse systematically the robustness of the results with respect to changes in the clustering methods, and (3) we investigate the effects of missing or partial temporal information on the detection of phases, as a number of models or biological data sets contain only temporal information on subsets of the proteins or genes of interest. Finally, we show that biological phases can also be inferred by using gene expression time series instead of protein-protein interaction data to build the temporal network. We make our code user-friendly and readily available for others to use for other biological systems (https://gitlab.com/habermann_lab/phasik).

## Results

### The Phasik workflow

The workflow of our analysis consists of two main steps: (i) building a temporal network representation of the cell cycle, and (ii) inferring cell cycle phases from it. This workflow, which we call Phasik, is illustrated in Figure 2.

**Figure 2:**
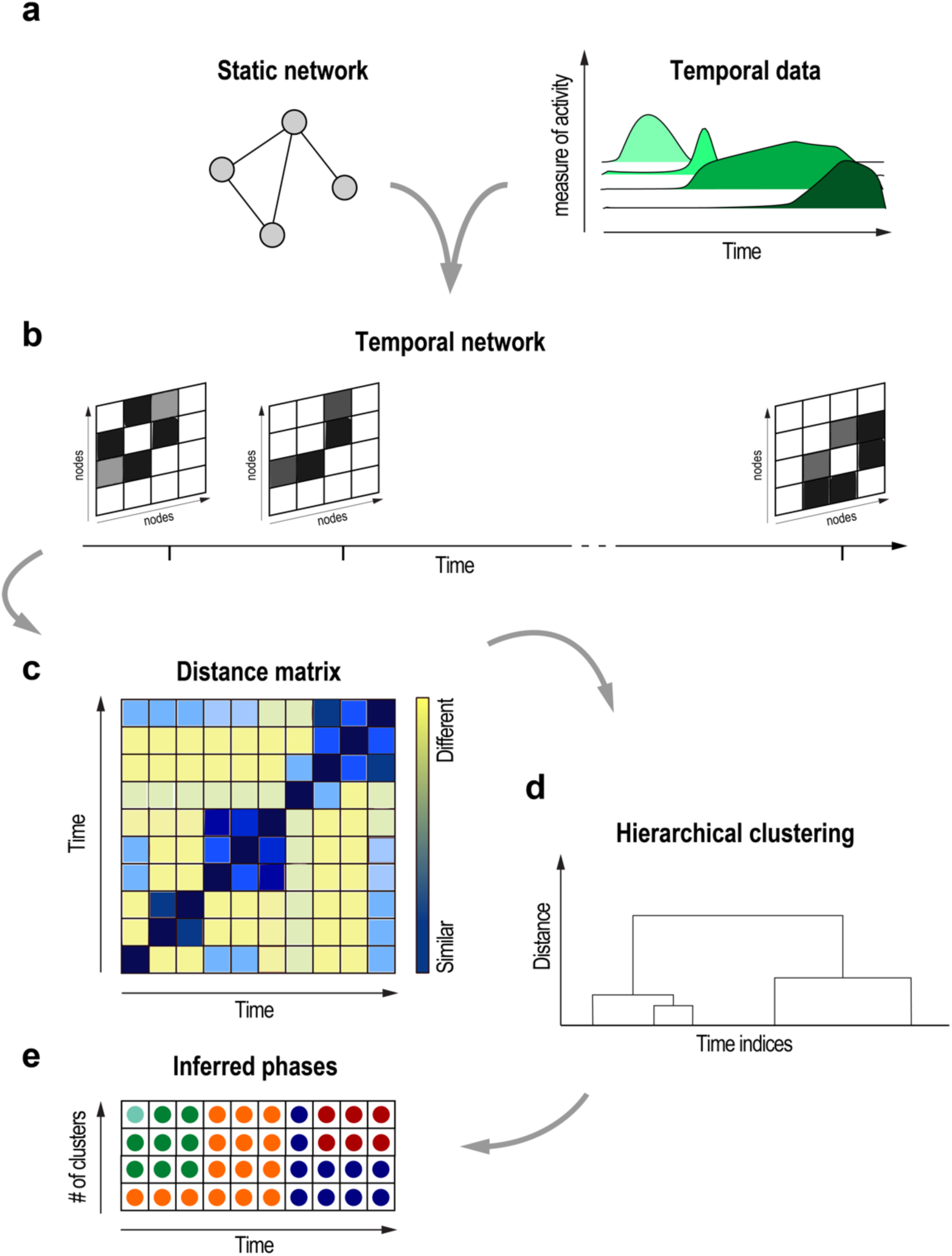
Multiscale phase inference with Phasik: schematic representation of the method. (a) Edge time series are integrated to a static PPI network to build (b) a partially temporal network, shown in snapshot representation. (c) Pairwise distances between snapshots are used to (d) cluster snapshots hierarchically. (e) The clusters obtained: each row corresponds to a fixed number of clusters. Each snapshot is shown as a dot whose colour represents the cluster it belongs to. Clusters can be interpreted as time intervals since there is a 1:1 correspondence between snapshots and timepoints. Each cluster can then be interpreted according to the underlying biological processes.

### Building a temporal network representation of the cell cycle

To build a temporal PPI network that represents the yeast cell cycle, we combined a static PPI network built from KEGG pathway data ((Kanehisa and Goto 2000), shown in Figure 3a), with temporal interaction data based on the mathematical model of the cell cycle as described by ((Chen et al. 2004), see Figure 3b). The static PPI network consists of 83 proteins (nodes) and 159 protein-protein interactions (edges). To obtain a temporal network, we first acquired time series of protein concentrations by numerical integration of the ODE model of (Chen et al. 2004), with a time step of 1 min over a full cycle of 101 min. For each edge A-B between proteins A and B in the static PPI network, we then defined its weight w_AB_(t) at time t as the product of the concentrations of the two proteins, in order to quantify their co-presence at that time (Wallach et al. 2013). Such a time-varying weight (see Figure 2a-b) could be defined only for the edges connecting two proteins which are described by the ODE model. In addition, the ODE model contains a few special variables representing the concentration of protein complexes: these quantities could also be directly used as evolving edge weights for the edges between the proteins forming these complexes (see Methods for details). Finally, we normalised each weight time series by its maximum value over the cell cycle, so that all weights vary between 0 and 1.

**Figure 3:**
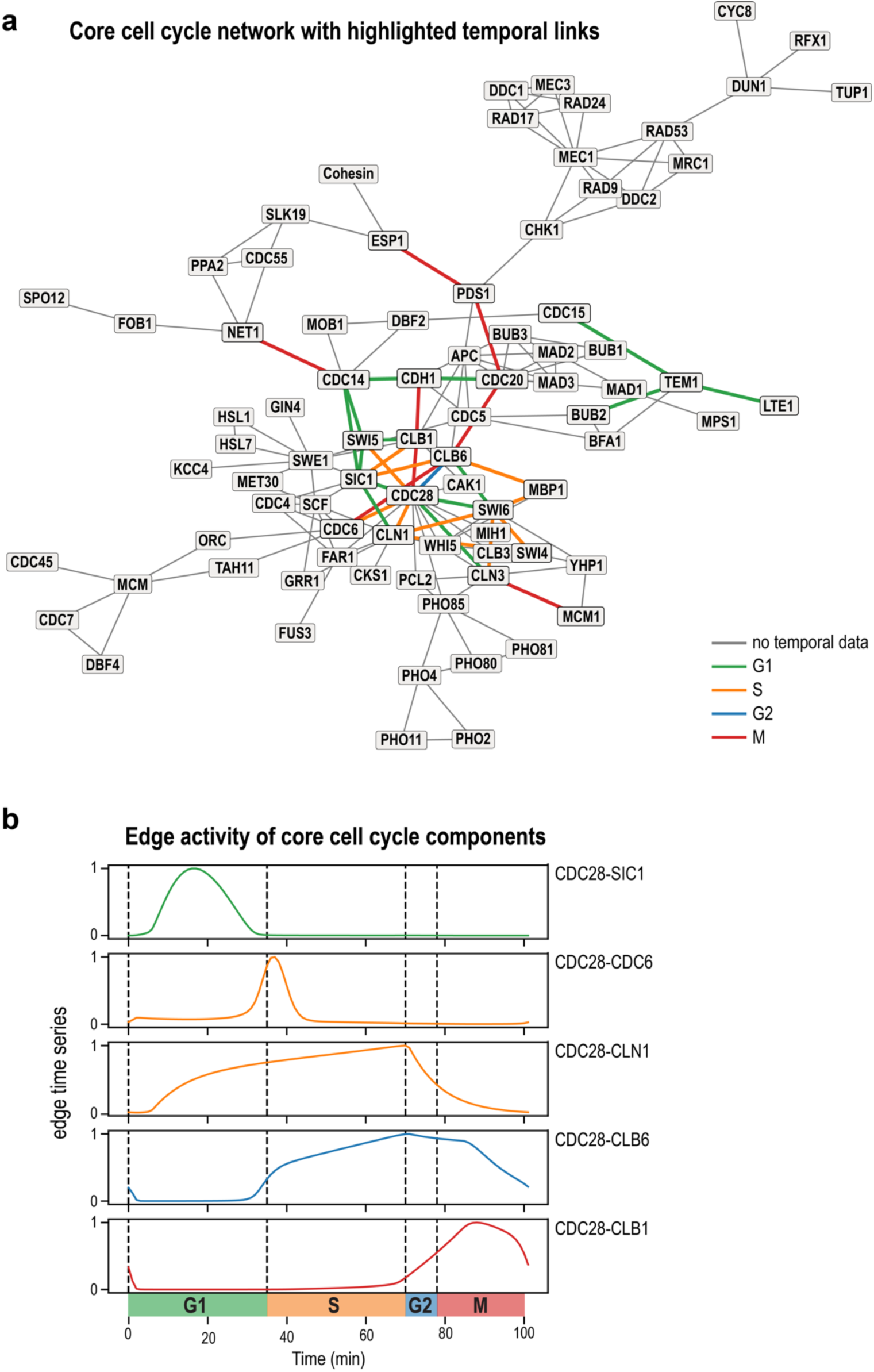
Partially temporal PPI network of the cell cycle. (a) Static representation of the temporal network: the initial static network contained 83 nodes and 159 edges. The 125 edges with no temporal information are shown in grey. The 34 temporal edges are shown in a colour that represents the phase of their peak activity. Colours are used for visualization only. (b) Edge times series: core cell cycle protein-protein interactions.

The ODE model provides overall temporal information only for a subset of nodes, and thus the edges, in the static PPI network, so that the resulting representation is a partially temporal network, containing both static edges for which no temporal information is available, and edges whose weights evolve in time (temporal edges). The weights of the static edges are set to a constant value of 1.

This procedure yielded time-varying weights for only 34 edges of the 159 edges (21%) of the PPI network. Several examples of the time series of edge weights obtained from the ODE model are shown in Fig. 3b (see Fig. S1 for all the time series), and the temporal edges are highlighted with colours in Fig. 3a (the colour representing the phase during which the weight is largest). Supplementary Movie S1 also shows an animation of the edge activity of the temporal cell cycle network.

Overall, the PPI temporal network can be seen as a series of n network snapshots (Figure 2b), each corresponding to a timestep at which the edge weights are observed. The minimal time between successive snapshots is equal to the temporal resolution used in the ODE numerical integration.

### Inferring phases from a temporal network

Since biological phases and processes are driven by specific protein-protein interactions, it is expected that each phase could be related to a specific structure or “state” (Masuda and Holme 2019; Pedreschi et al. 2020) of the temporal PPI network. In particular, a large similarity between snapshots of the temporal network at different times could indicate that the system is in the same phase at these times, and low similarity between successive times could indicate a change of phase (Masuda and Holme 2019; Gelardi et al. 2020; Pedreschi et al. 2020).

This idea can be taken further and formalised to infer phases by performing a clustering of the snapshots of the temporal PPI network. Such inference involves three steps (Masuda and Holme 2019), illustrated in Figures 2b-e: given a temporal network (Figure 2b), (i) we compute the distance between each pair of snapshots and build a distance matrix encoding all these distances (Figure 2c), (ii) using this distance matrix, we apply a (hierarchical) clustering algorithm on the snapshots (Figure 2d), and (iii) we extract the clusters, after choosing the desired number of clusters. Each cluster of snapshots can then be traced back to the set of timesteps corresponding to those snapshots and interpreted as a phase of the cycle (Figure 2e); see Methods for full details. Finally, the inferred phases can be compared to ground truth knowledge of the cell cycle and interpreted biologically. In most clustering algorithms, the number of clusters is fixed a priori. However, many real systems, including biological ones, exhibit dynamics on multiple timescales. Therefore, in order to explore the potential hierarchy of phases and subphases of the cell cycle, and to be able to infer phases across a range of timescales, we compute clusterings with varying numbers of clusters (Figure 2e): increasing the number of clusters potentially allows us to probe shorter timescales and identify subphases of the main cell cycle phases that are discovered at a small number of clusters. For example, computing 2 clusters gives us a more coarse-grained version of the temporal organisation than computing 5 clusters. For each number of clusters, we compute a quality metric of the clustering (the silhouette, see later) in order to investigate whether a preferred partition emerged from the analysis.

The phases determined by the clustering algorithm, at fixed number of clusters, may be affected by two main technical choices in the pipeline: (1) the distance measure used to compute the distance matrix between snapshots and (2) the specific clustering algorithm considered, together with its parameters (in particular, the way to measure a distance between clusters: while default settings exist in most numerical implementations of these algorithms, they can actually be changed and several choices are available). Unless stated otherwise, we used (1) the Euclidean distance and (2) hierarchical clustering with the “Ward” method of computing the distance between clusters in our analysis. These settings were chosen as reference because they provided the best clusters according to the following criteria: biological interpretability, quality of clustering, and robustness of the method.

We used the average silhouette score to assess the quality of each clustering (Rousseeuw 1987). This score is a number between −1 (the worst) and 1 (the best) that indicates how well separated the clusters are: values near 1 indicate a very good separation (hence a meaningful partition of the snapshots), while values near 0 indicate overlapping clusters. Negative values indicate that one of the snapshots was not correctly assigned to a cluster. In addition, we investigated quantitatively the robustness of the clusterings (at fixed number of clusters) with respect to the parameter choices by computing similarity scores between clusterings. To do this, we used the adjusted Rand index, which can take values between −1 and +1. A value of 1 indicates that two clusterings are identical (up to permutation of the labels) and a value of 0 indicates random cluster assignments. Negative values can occur when two clusterings are less similar than expected for random cluster assignments. See Methods for more details.

### Inferring the multiscale phases of the cell cycle

We inferred phases of the budding yeast cell cycle from the temporal cell cycle network, illustrated in Figure 3 and animated in supplementary Movie S1, using the Phasik workflow described above. First, we computed the distance matrix, as described above and shown in Figure 4a for snapshots observed at a temporal resolution of 1 minute. A visual inspection of this matrix revealed prominent dark blue diagonal blocks, corresponding to highly similar successive snapshots of the temporal network and thus indicating a marked temporal structure of the system. Second, we applied hierarchical clustering and computed clusterings with the number of clusters ranging from 2 to 11. The resulting partitions of the cell cycle timeline are shown in Figure 4b (Figure 4c displays the average silhouette that measures the quality of each clustering, as a function of the number of clusters). This quality score was around 0.5-0.6 in all cases, indicating that no partition should be discarded from this criterion. In fact, the roughly constant value of the silhouette rather indicates that many timescales are relevant in the system. The largest values of the average silhouette were obtained for numbers of clusters ranging from 5 to 9.

**Figure 4:**
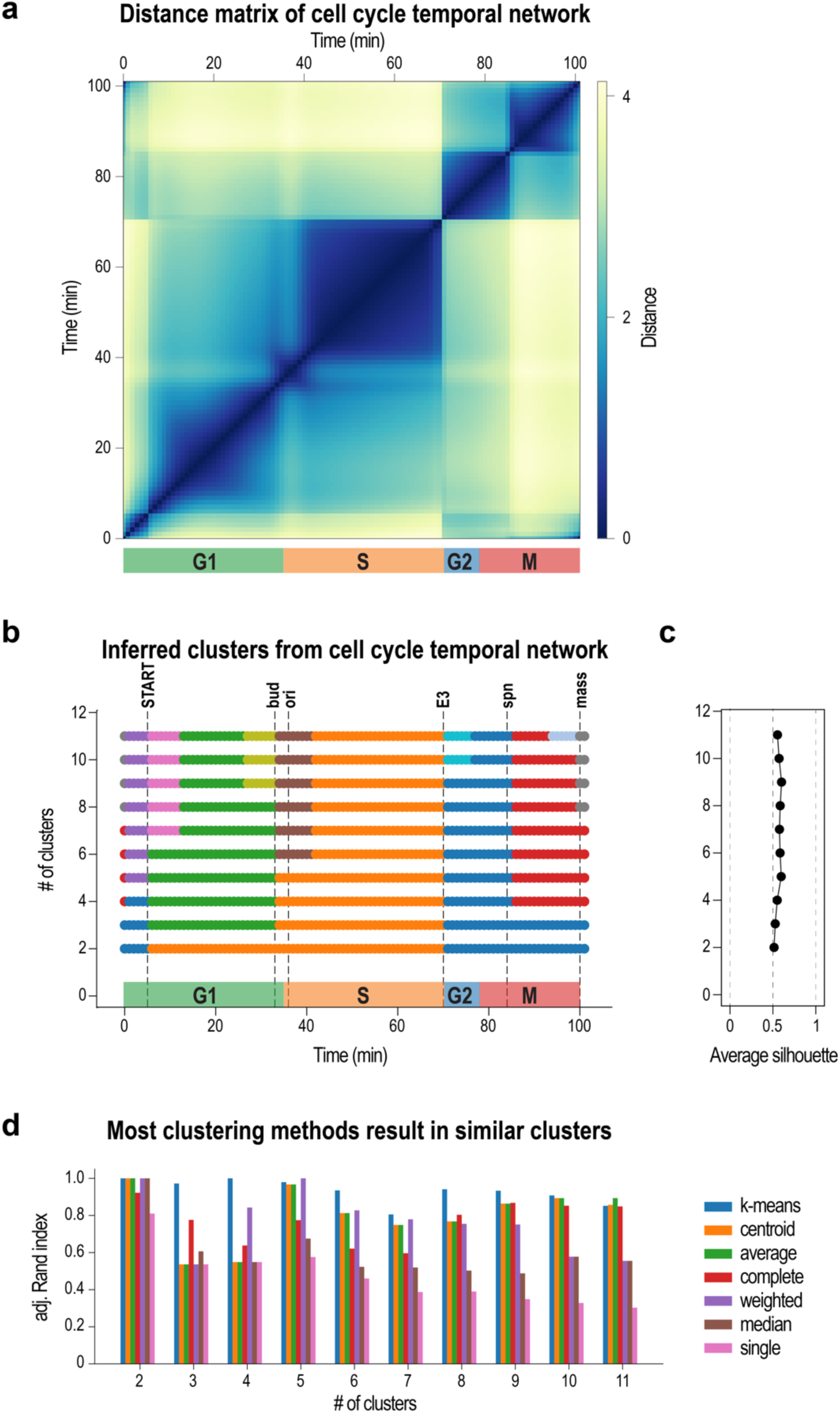
Cell cycle phases inferred for a range of timescales from the temporal PPI network shown in Figure 3a. (a) Distance matrix, using Euclidean distance. (b) Clusters inferred for a number of clusters ranging from 2 to 11. (c) Quality of clustering for clusters in (b), quantified by their average silhouette. Values close to 1 indicate well separated clusters. Similar values of about 0.6 indicate that no number of clusters can be readily discarded. (d) Most clustering methods resulted in similar clusters to the “Ward” method used in (b-c), as shown by the adjusted Rand indices.

To validate the results of the Phasik pipeline, we took advantage of the ground truth coming from biological knowledge, and checked that the starting and ending times of each cluster corresponded to the timings of known biological phases and events. We used the timings of the primary cell cycle phases G1, S, G2, M determined, relative to the budding yeast cell modelled in (Chen et al. 2004). In addition, we used the timings of the following checkpoints and physiological events as they were described in the same study: “bud” indicating bud emergence, “ori” the start of DNA synthesis, “spn” indicating completion of chromosome attachment to the spindle and alignment of chromosomes on the metaphase plate, and “mass” indicating the start/end of a cycle. Timings of the START checkpoint and the “E3” event (short excitation period associated with the sharp decrease in the SBF complex concentration) were determined by a stability analysis in (Lovrics et al. 2006).

Visual inspection of the distance matrix of the temporal cell cycle network, returned by Phasik, indicated 4 major clusters roughly corresponding to the G1, S, G2, and M phases of the cell cycle (see Figure 4a), as well as a small cluster at the very beginning of the cycle. The application of the clustering algorithm allowed us to perform a more detailed and precise analysis of the clusters, at each number of clusters (see Figure 4b) and to discuss them in relation with the evolution of the edge weights shown in Supplementary Figure S1:

- When using 2 clusters, the algorithm detected two phases: a phase corresponding to the G1/S phases, and one including the G2 and M phases mitosis. The detected G1/S phase starts at the START checkpoint, at which point the cell commits to entering the cell cycle. It ends at the E3 excitation peak before entering the G2/M cluster, which persists then until the START checkpoint. The main determinants for this clustering are the abrupt increase (START) and decrease (E3) of the activity of edges SWI4-SW6 and SWI6-MBP1. These two edges correspond molecularly to the activity of the SBF and MBF complexes, respectively, which are known to initiate at the START checkpoint. They are known to be activated by the CDC28-CLN3 complex early in the cycle and are required for the activation of gene expression of cyclins CLN1 and CLN2. The fact that these are the two first clusters to be detected highlights that our method is able to detect such abrupt changes in edge weights.
- For 3 clusters, the separation between phases G1 and S was detected at the physiological event of bud emergence (“bud”). Several molecular events can be linked to this separation: the emergence of the CDC28-CLB6 and SWI6-CLB6 edge weights which persist until the E3 event, as well as short peaks of the weights of CDC28-CDC6 and CDC6-CBL6. Moreover, the weight of CLB6-SIC1 decreases. Finally, the weights of CLN1-SIC1 and CDC28-SIC1 seem restricted to the G1 phase.
- At 4 clusters, the four phases G1, S, G2 and M were clearly detected, though the transition of the G2/M phases was obtained at the “spn” checkpoint. This checkpoint coincides with the abrupt change in edge weights for BUB2-TEM1, CDC15-TEM1, CDC20-CLB5, CDC20-PDS1, as well as LTE-TEM1. Biologically, it represents the moment in metaphase when all chromosomes should be properly attached in a bipolar fashion to the mitotic spindle. Once the spindle checkpoint is passed, the cell progresses into anaphase. Furthermore, the beginning of the G1 phase (before the START checkpoint) is clustered with the G2 phase, indicating that they seem to be more similar to each other than to other parts of the cell cycle. However, this split was not robustly found with other clustering methods, as is discussed in more detail below.
- At 5 clusters, the 4 phases G1, S, G2 and M were identified, with the G1 phase split in a pre-(purple in Figure 4b) and post-START (green) cluster. The cluster corresponding to phase G2 again extends to metaphase and the “spn” checkpoint.
- At 6 clusters, a small (brown) cluster appeared at the transition between phases G1 and S, immediately after the “bud” event, corresponding to a peak in CDC28-CDC6 and CLB6-CDC6 edge activities. CLB6 is known to be required for the initiation of DNA synthesis by activating CDC28, as well as the DNA-replication protein CDC6. Both interactions are thus initiating the transition from phase G1 to S.
- At 7 clusters, a new cluster was detected in phase G1, right after the START checkpoint (pink cluster). There are several changes in edge activities that can be linked to this new cluster: CDC14-SIC1 and SIC1-SWI4 have sharp activity peaks at this cluster. Furthermore, the edge activities of BUB2-TEM1, CDC14-CDH1, CDC14-SWI5, CDC15-TEM1, CDC20-CDH1 and CLB1-SIC1 drop during this phase and CDC14-NET1 shows a subtle increase. Finally, there is the transitions of MBP1-SWI6, SWI4-SWI6, CDC28-SIC1, CLN1-SWI4-CLN1-SWI6, CLN1-SIC1 from low to high activity that could contribute to this new cluster.
- At 8 clusters, a new cluster was introduced at the end of phase M, which extends to the beginning of the G1 phase (grey cluster). It can be linked to the late edge activities of CHD1-CLB1 and CDH1-SIC1: this is well in agreement with the fact that the APC-CDH1 complex is known to degrade CLB1 to induce mitotic exit. We can furthermore observe a drop in the edge activity of CLB1-SWI5 and CDC28-SWI5.
- At 9 clusters, a new (light green) cluster was found towards the end of G1 right before the “bud” event. The edge activity underlying this new cluster is the short and prominent peak in the edge activity of CLB6-SIC1 (Supplementary Figure S1). Biologically, at this stage, CLB5/6 are already prominently expressed and have formed stable complexes with CDC28. They are however, until the onset of phase S, inhibited by SIC1. Following the destruction of SIC1, replication can be initiated.
- At 10 clusters, the G2 phase was detected (light blue cluster), starting from the E3 excitation point. More subtle edge activity changes most likely contribute to this new cluster: the transition from high to low CDC28-CLN1, the subtle rise in CLB1-SIC1, the rise in CLB1-SWI5, CDC28-SWI5 and CDC28-CBL1 and the subtle increase of CLB6-SIC1.
- Finally, at 11 clusters, mitosis was further refined, with the detection of a separate anaphase (red cluster) and telophase (light grey cluster). This new split coincides with change in the activity of edges CDC14-NET1, CDC14-SWI5, CDC20-CLB6, CDC20-PDS1, CDC20-CLB1 and ESP1-PDS1.

Overall, the computed clusters corresponded thus to known biological processes across a range of timescales, and these clusters were of a high quality according to their silhouette scores. Note that, for high numbers of clusters, the clustering obtained presents much more finely time-resolved pictures of the temporal organisation of the cell cycle than with the major cell cycle phases, with subphases and biological processes happening over shorter time scales. A more detailed view of the silhouette scores is shown in Figure S2 for each case. For 6 or more clusters, negative silhouette values appeared for a few snapshots. As the average silhouette score was highest at 5 clusters and as the 5 clusters case seems to be the most biologically relevant for detecting the basic phases of the cell cycle, we use this case further in the rest of this study.

### Testing the robustness of Phasik against method variations

At each step of Phasik, several parameters can be set and methodological choices can be made. Each such choice can affect the resulting inference of cell cycle phases.

First, the definition of the evolving edge weights from the ODE model might affect the clustering results. To investigate this further, we built a temporal network with the same method as above, but without normalising the weights by their maximum value. The resulting distance matrix (see Supplementary Figure S3a) and clusterings (Supplementary Figure S3b) differed from those obtained using normalised weights (shown in Figure 4). Indeed, the G1/S phases were less well defined. The “bud” and “ori” events were not detected. “START” was only detected when using at least 8 clusters and the G1/S phase boundary was shifted. The G2 and M phases were found up to 5 clusters, as with normalised weights. A shorter G2 phase was detected already at 7 clusters and the period between “E3” and “spn” was further split with higher cluster numbers. These differences are likely due to the large differences in concentration values found in the ODE: they indeed vary between 1 and 200 units. Protein interactions resulting from very low concentrations therefore lead to small edge weights and their variations influence clustering much less than larger weights, even if they might be biologically equally important. It seems thus important to use a normalisation step when defining the edge weights in the temporal network from which one aims to infer relevant phases.

In the second step of Phasik, a distance metric needs to be chosen to compute the distances between snapshots (Masuda and Holme 2019). Moreover, a clustering algorithm needs to be selected (e.g. hierarchical or k-means), together with its associated parameters (Figure 2d).

To test the robustness of the Phasik workflow against variations in the methods and parameters chosen, we compared different clustering methods and different clustering parameters to our base method (hierarchical clustering with Ward linkage). Specifically, we used hierarchical clustering with the following 6 linkage methods: median, centroid, weighted, average, complete, and single (see (Müllner 2011) for details), as well as k-means clustering, which is non-hierarchical. For each of these methods, we computed clusterings and quantified how similar they were to those of our base method, using the adjusted Rand index. We show the adjusted Rand index for all 7 methods and for each number of clusters in Figure 4d. Actual clusters for each case are shown in Supplementary Figure S5, together with average silhouettes. As shown in Figure 4d, the clustering was most robust when using 2 clusters: most methods found identical clusters, except hierarchical clustering with complete and single linkage (which were still very similar with an adjusted Rand index close to 1). For 3 and 4 clusters, k-means yielded clusters almost identical to Ward, but the other methods gave fairly different clusters, except for complete and weighted, which showed slightly higher adjusted Rand indices. At 5 clusters, most methods yielded results very similar to Ward, except for median and single with adjusted Rand indices close to 0.6. For larger numbers of clusters (6 and more), the various methods still agreed fairly well with Ward, especially k-means and, to a lesser extent, hierarchical clustering with average, as well as complete linkage. The methods that performed most differently were median and single, with adjusted Rand index values close to 0.4. Remarkably, “k-means” performed overall most similarly to the hierarchical clustering with “Ward”, even though it is not hierarchical. This could indicate an inherent hierarchy in the temporal structure of the system.

By definition, different clustering methods are expected to favour different shapes of clusters, and hence yield different clusterings. Identifying similar clusterings with different methods therefore is an indication that the underlying data contains strong phasic information. In addition, we also investigated the robustness of the method against changes in the distance metric used (see Supplementary Figures S5). We compared Euclidean distance to 3 other distance metrics: cosine, Manhattan, and Chebyshev. All distance metrics give nearly identical clusterings at each number of clusters. Only Chebyshev at 2 clusters yields a Rand index smaller than 0.6.

### How much temporal information is needed to infer meaningful phases?

The framework of temporal networks is usually considered when temporal information is available for all pairs of nodes: the data is a set of temporally resolved interactions, and all edges are then temporal. Here instead, the temporal information is available only for a fraction (21%) of the edges (protein pairs), and we have for simplicity set a constant weight to the other edges of the PPI. Such a situation of partial temporal information might in fact happen in many systems, in particular in biological systems, where experimental measurements of temporal information about all interactions are currently not feasible. We thus investigated how the detection of meaningful phases through the Phasik workflow depends on the amount of temporal information available. To this end, we artificially discarded the temporal information from selected edges, building in this way modified temporal networks. We then computed clusters from these networks and compared them to those obtained with the original temporal network constructed with all the available temporal information.

First, we discarded temporal edge activity for each specific edge by setting its weight to a constant value (e.g., 1) and computing the partition of the snapshots into 5 clusters using Phasik. We repeated this procedure for all temporal edges. As shown in Supplementary Figure S6, discarding the temporal information of one single edge resulted in nearly identical clusters in most cases, indicating the robustness of the clustering against small alterations in edge weights. Small differences occur only in a few cases: for instance, removing the variation of the temporal edge weight representing the interaction between CDC28 and CLN1 resulted in a slightly earlier onset of the S phase compared to the original temporal cell cycle network (Supplementary Figure S6 c, d). Supplementary Figure S6e also highlights (top of the plot) several other edges, where variations in the clustering were observed when their weight variations were ignored.

Second, we focused on nodes rather than single edges, to investigate whether some nodes are more central than others (i.e. more important for extracting meaningful phases). In other words, we studied whether the temporal information of a specific node and its interaction partners was more informative than for other nodes in terms of correctly detecting sensible phases. This question is especially meaningful since edge weights were defined based on time series related to nodes. We thus built temporal networks using temporal information restricted to one specific node and its interaction partners, setting all the other edge weights to a constant value. We then computed the clusters from the Phasik pipeline with 5 clusters and compared the result to the one of the whole temporal network using the adjusted Rand index. We performed this procedure for each node present in the ODE model, respectively. Figure 5a shows the adjusted Rand index obtained in each case. In some cases (e.g. CLB6, CDC28 or CLN3), the similarity with the clustering obtained with full temporal information remained quite large. For some other nodes however, very different clusters were obtained (e.g. CHD1, CDC14, CDC6, see Figure 5a). Figures 5b-e and S7 show more details of the distance matrices and clusters obtained in various cases. When temporal information was restricted to the edges connected to CDC28, the original 5 clusters were reproduced fairly well (Figure 5b,c). This is not unexpected, as CDC28 has 8 temporal edges and plays a central role in regulating the cell cycle. Notably, a protein like MBP1, which has only to 2 edges with temporal information, also yielded clusters quite similar to the original ones (Figure 5d,e): when keeping only temporal edge weights for MBP1, the clustering detected G1 more reliably than the other phases, while the S phase was not properly clustered and split into three phases. This result can be explained by the temporal profile of MBP1 and its partners (see Supplementary Figure S1), with constant activity over G1 and S of MBP1-SWI6 and varying activity of CLB6-MBP1.

**Figure 5:**
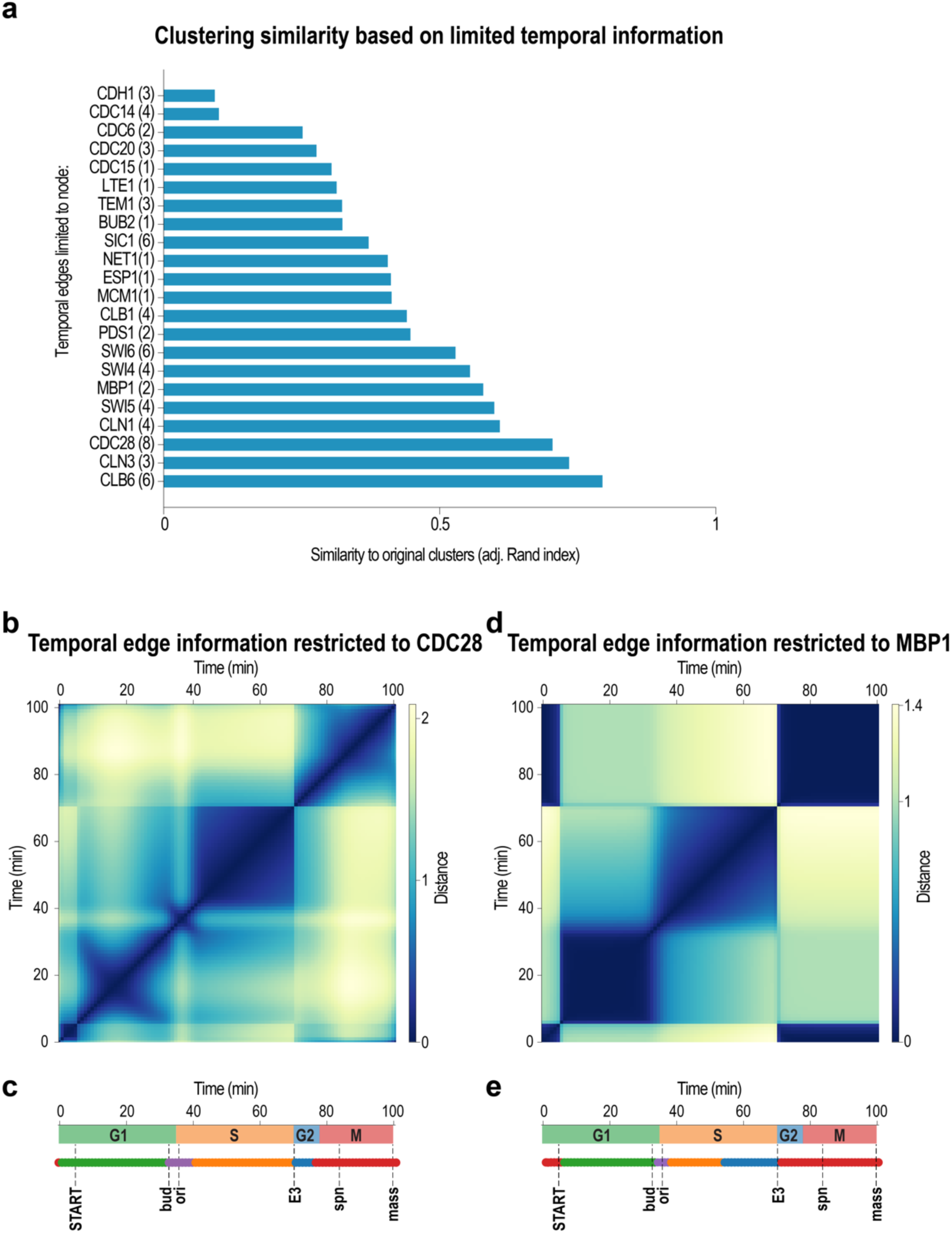
Only a few selected temporal edges were sufficient to recover the original 5 clusters of Figure 4. Here, we kept only the temporal information of edges from a given node, and set all other edge weights to a constant value. (a) Similarity of the newly computed 5 clusters to the original 5 clusters, for each temporal node (with its number of temporal edges in parentheses). Values closer to 1 indicate clusters close to the original ones. Examples of (b),(c) CDC28 and (d),(e) MBP1, which have 8 and 2 temporal edges, respectively. (b) and (d) show distance matrices and (c) and (e) show the 5 computed clusters (bottom coloured lines).

Examples of nodes whose sole temporal information recovered the original 5 clusters less well are shown in Supplementary Figure S7. Temporal edges of CLB1 detected the G2 and M phases better, but were unable to detect the G1 and S phases (Supplementary Figure S7a). This agrees with the activity profile of CLB1 edges: CLB1 has 4 temporal edges, but they are all active at the very beginning of the G1 phase only, as well as prominently in phases G2 and M (Supplementary Figure S1). Temporal information on SIC1, on the other hand, detected only one cluster over the S, G2, and M phases, whereas the G1 phase was split into 3 distinct clusters. The edge weight timelines of SIC1 and its partners (6 active edges) are largely restricted to the G1 phase, with only CLB1-SIC1 being active late in the M phase as well (Supplementary Figure S1).

We finally investigated whether some features of the nodes could predict their importance in this procedure, as determined by their possibility to recover clusters similar to the original ones if only their activity timeline and the ones of their partners are known (Supplementary Figure S7e). We found only very weak correlations between the adjusted Rand indices of Figure 5 and the number of edges with temporal information with which a node participates. Similarly, only very weak correlations were found with typical measures of centrality for static networks such as degree or betweenness centrality, with several nodes having low centrality but high adjusted Rand index. Only the eigenvector centrality showed a slightly larger (but still weak) correlation with the adjusted Rand index.

### Influence of sampling frequency of edge time series on clusterings

Another issue related to the amount of temporal information needed concerns the temporal resolution of the evolving weights, corresponding to the inverse of the sampling frequency. While we have used until now the finest possible resolution (i.e., the temporal resolution with which the ODEs have been numerically integrated), concentration time series might a priori be available only at coarser resolutions. We thus downsampled the original time series for a range of temporal resolutions to determine the temporal resolution needed to infer meaningful phases. The resulting distance matrices between snapshots are shown in Figure 6 for snapshots observed every 5 or 15 minutes, together with the 5 clusters obtained by the clustering algorithm. Despite the coarser resolution, results showed good agreement between clusters and known phases (Figure 6). With a temporal resolution of 5 minutes, all 5 clusters were robustly obtained (Figure 6 a,b). Even with a temporal resolution of 15 minutes, the original clusters could reliably be obtained (Figure 6 c,d). Clusters are shown for temporal resolutions ranging from 2 to 20 minutes in Supplementary Figure S8. Note that average silhouette values became lower with coarser temporal resolution. Furthermore, lower sampling frequencies yielded fewer timepoints and, consequently, fewer clusters could be computed: a phase could be detected only if it was represented by at least one timepoint. Combined, these effects imply a lower bound on acceptable sampling frequencies that will depend on the timescales of interest in the system considered.

**Figure 6:**
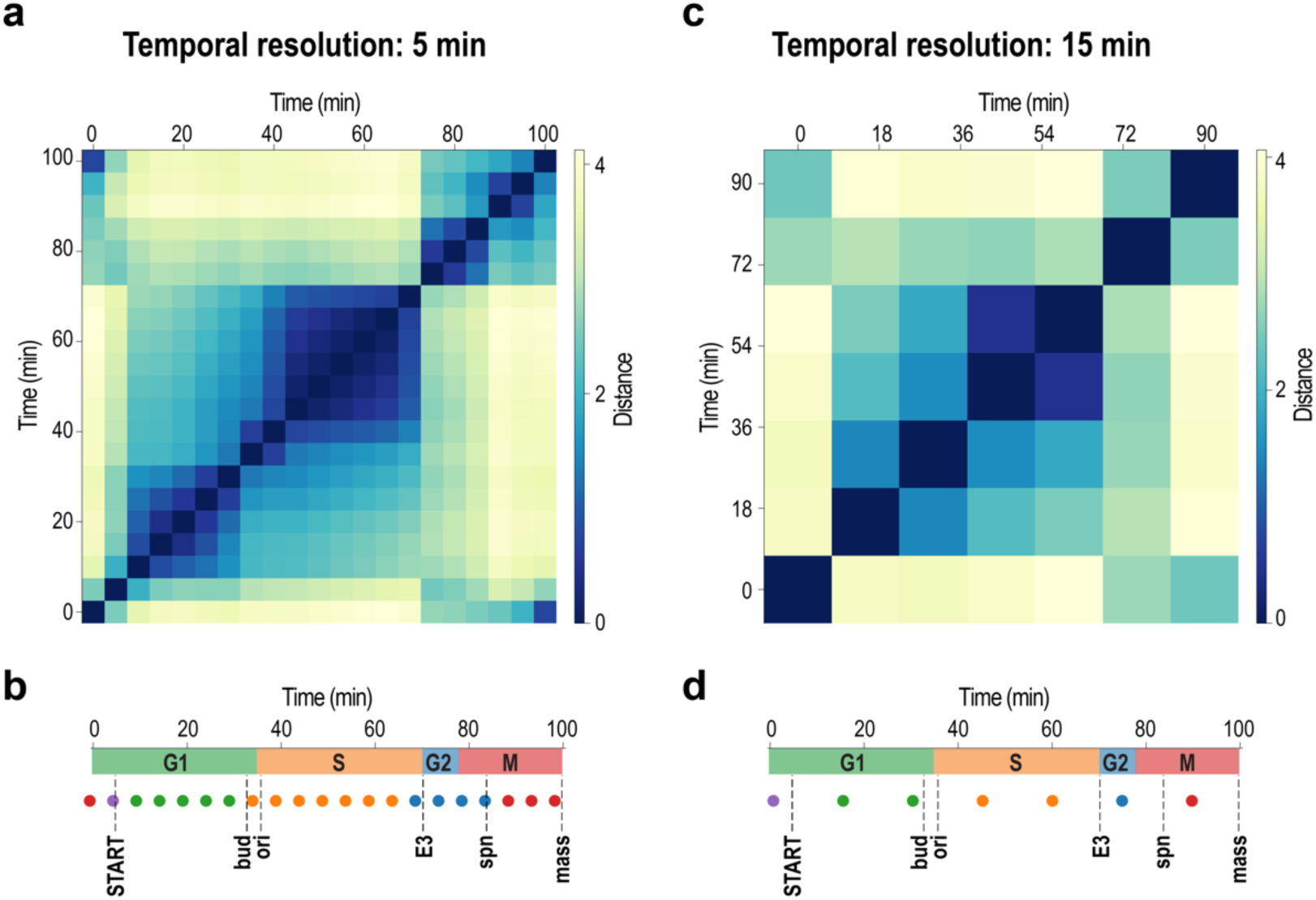
Biological phases inferred with downsampled time series. (a) and (c) use timepoints every 5 min, (b) and (d) every 15 min. The original timestep was 1 min. The clusters, shown as lines of coloured dots, still detected the main biological phases. These results show that 15 min is the minimum timestep needed to detect shorter phases.

### Inferring cell cycle phases from gene expression data

The previous results were obtained using the protein time series obtained from the ODE model by Chen et al. (Chen et al. 2004). These time series have the advantage of being noise-free and having virtually infinite sampling frequency (as one can numerically integrate the ODE with an arbitrary small timestep). However, such high-quality mathematical models are so far only available for a few biological systems and are mostly restricted to systems with sufficient biological knowledge. It is therefore important to investigate whether cell cycle phases could be inferred in a similar way from gene expression time series data emanating from RNA-sequencing or microarray studies. Such data is indeed much more easily available and provides temporal information for virtually all genes. To test this, we used a time-series RNA-sequencing dataset described in (Kelliher, et al., 2016): for this study, yeast cells had been first synchronised, then were released to undergo three full cell cycles, with RNA samples taken from the culture every 5 minutes for RNA-sequencing. In the published dataset, RNA-levels of all genes are available. One full cell cycle was reported to last 75 minutes. For our study, we only used the first cell cycle, as the time series deteriorate due to de-synchronisation of the cells. We downloaded normalised read counts and then normalised each time series so that values were between 0 and 1. As previously, we used the same static PPI network from KEGG, now deriving edge weights from gene expression data rather than protein concentrations of the nodes, to build a temporal network. Then, we inferred phases, following the workflow described above. The resulting temporal network contained 83 nodes and 159 edges, of which 158 contained temporal information over 16 timepoints.

Visual inspection of the distance matrix indicated a less clear temporal structure (Figure 7a) than that of the temporal network built from protein concentrations. This could be due to a higher noise level in the data, as well as less clear transitions in gene expression from one state to the next. We next inferred clusters using Phasik (Figure 7b). We identified 4 stable clusters that roughly correspond to the 4 cell cycle phases and that persist at higher numbers of clusters. The average silhouette values were lower than the ones reported when using the temporal PPI derived from the ODE model, indicating a worse separation of clusters. Values ranged from 0.25-0.3, with a maximum just above 0.3 at 5 clusters, and values below 0.25 for 8 clusters or more (Figure 7c).

**Figure 7:**
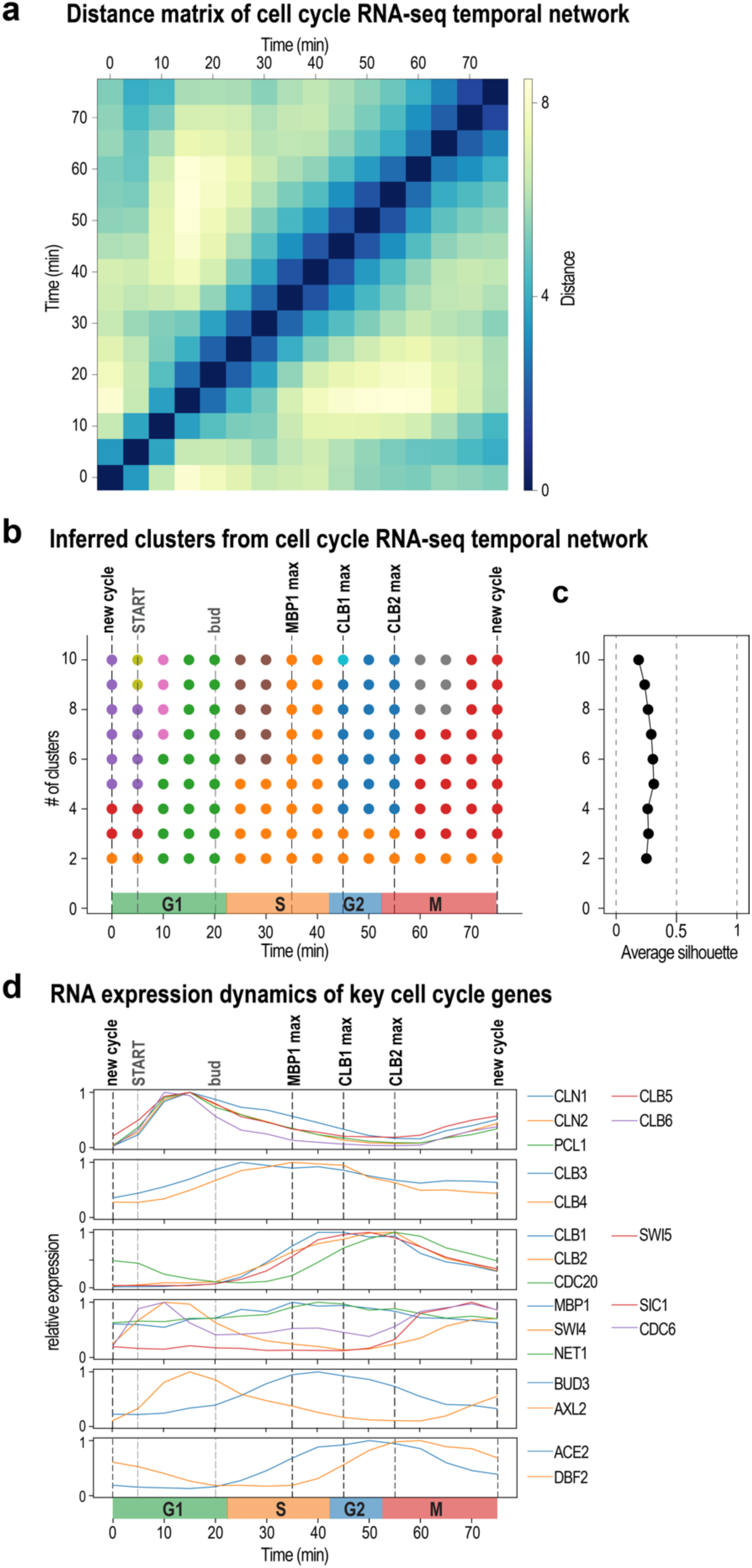
Phase inference from a temporal network constructed using gene expression data from RNA-sequencing. (a) The distance matrix reflects the general coarser resolution of the data. (b) inferred phases were similar to the ones inferred from protein concentration data obtained by the ODE model and could identify the 4 phases of the cell cycle robustly. (c) Average silhouette values were low, indicating lower quality of the clustering. (d) Dynamics of RNA expression of key cell cycle genes.

Interpreting the resulting cell cycle phases from the temporal RNA-seq expression data was much more challenging than from those inferred from protein concentration data from the Chen model. We mainly used evidence of temporal profiles of cell cycle genes known to be temporally regulated in the cell cycle from to assign cell cycle phases (see reviews from (Nasmyth 1993) and (Koch and Nasmyth 1994); with additional data from (Lord et al. 2000); and (Amoussouvi et al. 2018)) (Figure 7d). The phases we inferred from the clusters fitted reasonably well to the G1, S, G2 and M phases, especially for 4 and 5 clusters. Edge weights of the most active edges for 4 and 5 clusters are shown in Supplementary Table S1. At higher numbers of clusters, we obtained new clusters that were similar to those obtained from the temporal PPI network, namely: a split of the M phase cluster with the pre-START G1 phase cluster; a separation of phase S in an early and a later phase; a separation of the G1 phase in early and late, as well as the split of the M phase in two separate subphases.

## Discussion

In this study, we inferred biological phases of the budding yeast cell cycle by representing it as a temporal network of protein-protein interactions and applying a clustering algorithm to the resulting series of snapshot networks. To build the temporal network, we started from a static PPI network in which we assigned time-evolving weights to certain edges, obtained by integrating time series of protein concentrations from an ODE model, as well as from expression data from an RNA-seq study. We inferred biological phases from the temporal PPI network for a range of potential timescales, by using the clustering algorithm with various numbers of clusters. We tested our method extensively against variations in algorithms and parameters chosen. We showed that our method is able to infer meaningful biological phases, corresponding to the principal cell cycle phases G1, S, G2 and M, with a finer resolution in subphases at higher numbers of clusters. The identification of these cell cycle phases was possible with both protein concentration data and gene expression data. We showed that our approach is robust against changes in distance metrics and clustering methods and identified those that performed best for our system of interest. It should be noted, however, that the clustering algorithms could perform differently for other types of data. A careful evaluation of different methods would therefore be of interest for other systems.

At each number of clusters, we matched the inferred phases to actual biological phases and processes and found good agreement with published data on the cell cycle. This may not be surprising as the mathematical model developed by Chen et al. is based on extensive knowledge of key cell cycle events. It is remarkable, however, that we obtained these results even though our network is only partially temporal, with temporal information missing for nearly 80% of edges. We stress here that no additional information about the cell cycle phases was used to infer these phases through the clustering method: results shown here are based only on the temporal network itself.

A peculiarity of the system we have considered is precisely that only partial temporal information was available. Usual temporal networks study indeed consider all edges to include temporal information (either discretised or with evolving weights). Here, we further investigated this point and the specific role of the temporal information in the inference of biologically meaningful phases. First, we showed that the inferred clusters remained almost unchanged if we discarded the temporal information of any single edge. Second, we showed that meaningful clusters could still be obtained even when temporal information was included for the edges of some single nodes. Results depended on the node selected, with some nodes carrying more important temporal information than others (i.e., being in some sense “temporally central”). For example, MBP1 (participating in 2 edges with temporal information) and CDC28 (participating in 8 such edges) yielded results comparable to the ones obtained with the original network, but CDC14 (participating in 2 temporal edges) did not. Interestingly, we did not find any correlation between the number of temporal edges of a node and its ability to recover the 5 cell cycle clusters. Furthermore, no correlation with typical static centrality measures could be observed, including degree, betweenness centrality, and only weak correlated was observed with eigenvector centrality. Third, we showed that a minimum sampling frequency was dictated by the timescales of the phases that should be detected, but that sampling frequency had minor effects on clusters at values above that threshold. Many open questions remain of interest to both theoreticians and experimentalists concerning partial temporal networks. Indeed, by understanding which interactions are most informative about the temporal structure of a system, experimentalists could focus primarily on those. For instance, while it is well-known in the cell cycle that CDC28 is a major driver of cell cycle events, major drivers might be less well known in other biological systems. Yet, our method shows that obtaining even partial temporal information on some components might be sufficient to gain insights into the temporal structure of a biological process.

To test the Phasik workflow, we have used a well-defined time series of protein concentrations obtained from an ODE model. This data has virtually infinite sampling frequency and is free of noise. Moreover, such models typically contain just a handful of proteins that were carefully selected to reproduce the dynamics of a specific biological system. However, such complete and high-quality models are not available for many biological systems, as they require a large body of experimental evidence and modelling effort. More accessible temporal data can be obtained by high-throughput measurements of RNA (RNA-sequencing, microarrays) or protein levels (quantitative proteomics) in time-course experiments. Such studies have become standard and are technically easier. However, high-throughput measurements come with downsides: 1) inherent high noise levels; 2) coarse temporal resolution, typically in the range of hours rather than minutes; 3) in the case of RNA-sequencing, transitions in RNA expression levels from one state to the next are much less marked than for proteins. We used RNA-sequencing data from a yeast cell cycle time series experiment to test whether we could infer biologically relevant phases with our method. Even though the data suffered from all the issues discussed above, we succeeded in inferring the 4 relevant cell cycle phases. Thus, while ODE models have many advantages over time series of expression data for the present application, the latter is a more available option for biological systems in general and can be readily analysed using the Phasik workflow.

## Conclusions

In conclusion, we inferred phases and subphases of the cell cycle by modelling it as a temporal network of protein-protein interactions, and clustering its snapshots. Additionally, we systematically tested the robustness of the results. We showed that similar results are obtained using protein concentration time series and gene expression data from RNA-seq. Finally, we investigated the effect of having a partially temporal network. While we focused on the cell cycle of budding yeast, many other biological systems share its features of interest for the study: time-varying interactions, partial temporal information, and a temporal structure that can be divided into phases. For this reason, this framework is very general and could be applied to many other biological networks. The method could also be used to predict phases in less known biological systems. We have made our code available online for interested researchers to use on other biological systems of interest (https://gitlab.com/habermann_lab/phasik).

## Methods

### Building the temporal network: integrating temporal information to a static network

A temporal network is a network where the edges connecting the nodes can vary over time. In a weighted network, this means that the edge weights are time-varying. We build such a (weighted) temporal network by incorporating time series of edge weights, which represent the activity of the corresponding interactions, into a static network. Note that we only consider undirected interactions.

In the case of proteins, actual protein-protein interactions are difficult to measure over time. Quantities relative to each protein are more accessible however, such as protein concentrations or associated gene expression levels. We used these (node) time series to generate corresponding edge time series. We multiplied the time series relative to protein A by that of protein B to obtain a time series for the interaction A-B. We did so only if this edge A-B exists in the static network. It is also possible that no edge time series for an edge A-B in the static network exists: this can happen if there are no time series for proteins A or B. In these cases, we set the weight to a constant value.

Our code to build temporal networks as described is available online (https://gitlab.com/habermann_lab/phasik). Several utility functions are available to easily integrate node or edge times series, e.g. RNA-seq data, to a static network. Details about the actual data used is provided below.

### Inferring biological phases from a temporal PPI network by clustering snapshots

A temporal network can be seen as a list of snapshots, where each snapshot is the adjacency matrix of the network at a given time. We expect two snapshots S and S’ at times t and t’ to be similar if the system is in the same state or phase at those times. On the contrary, if times t and t’ correspond to different phases, we expect the snapshots at those times to be very different. The underlying assumption is that the structure of the network is linked to the state that the system is in.

This idea can be formalised by clustering the snapshots of a given temporal network (Masuda and Holme 2019). Each snapshot is a data point and is assigned to a cluster. As an output, we obtain clusters composed of snapshots: if snapshots S and S’ are in the same cluster, it means that the system is in the same phase at times t and t’. We mainly used hierarchical clustering and k-means clustering. These were implemented with functions from Scipy (Virtanen et al. 2020) and Scikit-learn (Pedregosa et al. 2011), respectively. To compute the distance between snapshots, we flattened each adjacency matrix and then used Euclidean distance as a vector distance metric, unless stated otherwise. To compute the distance between clusters in the hierarchical clustering, we used the Ward variance minimisation method, unless stated otherwise. The desired number of clusters was set a priori, both in the hierarchical clustering used and in k-means.

### Measures of clustering quality

To check the quality of computed clusters, we used the silhouette score (Rousseeuw 1987), which ranges from −1 to 1. For a given data point, the silhouette score is larger if it is close to other data points in the same cluster (cohesion) but far from data points in other clusters (separation). The average silhouette score is obtained by averaging over all data points and is a measure of how well separated the computed clusters are. Low scores indicate clusters of bad quality. To compare clusterings, we used the adjusted Rand index which ranges from 0 to 1. Its value is 1 when the two sets of clusters are identical, but close to 0 for random cluster assignments.

### Building of static PPI network of the cell cycle from KEGG

We built a static PPI network of the budding yeast cycle from the manually curated regulatory network downloaded from the KEGG database (Kanehisa and Goto 2000). We built our PPI in two main steps: we first merged duplicate nodes and then converted nodes and edges of multiple types to nodes of a single type connected by undirected edges of a single type. The final PPI network, shown in Fig. 2(a), consists of 83 proteins (nodes) and 159 protein-protein interactions (edges). Note that both, protein expression, as well as RNA-seq expression data can be mapped to this network. The nodes thus represent both, proteins, as well as genes.

In the KEGG network, nodes can be of 5 types: gene, group, compound, map, and ortholog. First, we discarded the 2 compound, 4 map, and 3 ortholog nodes which do not represent genes or proteins, and kept only the 125 gene nodes (e.g. CDC28) and the 27 group nodes (e.g. [CDC28, CLN3], representing sets of gene nodes). We then merged duplicate nodes and removed disconnected nodes. For group nodes, we distinguished true groups from normal groups: true groups are node groups with components only interacting among themselves except for one edge that connects them to external nodes.

This distinction allowed us to treat true groups as one node (e.g. nodes [SMC1, SMC3, MCD1, IRR1] in the PPI network were replaced by a new node, Cohesin). This resulted in 83 nodes in the static PPI network. Second, we converted the 88 KEGG relations to edges in the PPI network. KEGG relations are of 3 types: gene-gene, gene-group, and group-group. For gene-gene relations, we simply added an edge between the corresponding nodes in the PPI network. For relations involving groups, we treated them as gene-gene in case of true groups. In the case of normal groups, we added an edge between each node of the group and the external node or nodes. Finally, we added edges between each pair of nodes that are part of the same normal group, and removed self-edges in the PPI network. This resulted in the 159 edges in the PPI network.

### Times series data for edge activities

We used two different datasets and built two different temporal networks by integrating them to the static PPI network presented above.

### Protein concentrations

We used time series of protein concentrations obtained from the reference ODE model of the budding yeast cell cycle (Chen et al. 2004). This mathematical model, based on and validated by extensive experimental data, describes the evolution of the concentration of a selected number of proteins over the entire cell cycle, based on knowledge about their interactions. We obtained the time series of protein concentrations by simulating the ODE model from the publicly available script of (Chen et al. 2004) with a time step of 1 min. The cell cycle in that study lasts 101 min, and hence there are 101 timepoints for one full cycle. The model consists of 46 variables that can be divided into 3 types: proteins (e.g. CDC6), protein complexes (e.g. the variable C2 represents the complex CLB1-SIC1), and 4 special variables (MASS, BUD, ORI, SPN) that serve as indicators of specific events along the cycle. Note that some proteins in the static PPI network are not represented in the ODE model, and vice versa. In addition, similar cyclins are represented by a single variable in the model, hence, we used CLN1, CLB1, CLB3, and CLB6 to represent CLN1/2, CLB1/2, CLB3/4, and CLB5/6, respectively.

For proteins A and B in the model, we defined the activity w_A-B_(t) of their interaction at time t as the product of their respective concentrations: w_A-B_(t) = [A](t) x [B](t), if there is an edge A-B in the static PPI network. Some edges in the PPI network are represented by a single “protein complex” variable in the model: in those cases, we defined the activity of that edge as the concentration of that single variable. We then normalised each of these times series.

### Gene expression from RNA-sequencing data

To create a temporal cell cycle network based on RNA-sequencing data, we used mRNA counts of a gene expression time series, normalised to the library size, from (Kelliher et al. 2016). We only considered the 15 timepoints of the first full cell cycle, starting at minute 10 in the dataset.

We defined the activity of a protein interaction A-B over time by the product of the respective RNA counts of the corresponding genes, as above: *w*_A-B_(t) = N_A_(t) x N_B_(t). We then normalised each of these times series.

### Temporal networks of PPI

We built two (partially) temporal networks of protein interactions for one full cell cycle of budding yeast: for both, we used the static PPI network described above, to which we integrated the edge activities from (i) protein concentrations, and (ii) the gene expression data from RNA-sequencing. Both contained 83 nodes and 159 edges, but a different number of temporal edges. In both cases, we set the weight of the other edges that lack temporal information to a constant value of 1.

### Data availability

The static protein interaction data used to build the static PPI network comes from the KEGG database, entry sce04111. The ODE model study of the cell cycle can be found in (Chen et al. 2004), and the authors have made their simulation script freely available at http://mpf.biol.vt.edu/research/budding_yeast_model/pp/getwinpp_current_model.php. The RNA-sequencing gene expression data was published in (Kelliher et al. 2016) and is available from GEO under accession number GSE80474. The code produced for this study is available in the form of a python package (Phasik) and notebooks that use the package at https://gitlab.com/habermann_lab/phasik.

## Acknowledgments

We want to thank Stefan Heldt and Béla Novàk for early discussions of their model. We thank the members of the Habermann team for critical reading of the manuscript. The project leading to this publication has received funding from the « Investissements d’Avenir » French Government program managed by the French National Research Agency (ANR-16-CONV-0001) and from Excellence Initiative of Aix-Marseille University - A*MIDEX. AB acknowledges partial support from the ANR project DATAREDUX (ANR-19-CE46-0008-01).

## Author contributions

ML, BHH, LT and AB conceived the study, ML was the main responsible for code implementation and data analysis, with help from AM and AT. ML, BHH, LT and AB were responsible for data interpretation. ML and BHH wrote the manuscript, with input from LT, AB, AM and AT.

## Description of Supplementary Data

**Supplementary Figures S1 - S8 contain data supporting main Figures 1 - 7.**

**Supplementary Movie S1:** animation of temporal edge activity in the cell cycle network based on the data from Chen et al., (Chen et al., 2014).

**Supplementary Table S1:** all active edges (weight > 0.7) for the temporal cell cycle network for 4 identified phases based on time series RNA-sequencing data (Kelliher, et al., 2016).

## Supplementary Figures

**Supplementary Figure S1:**
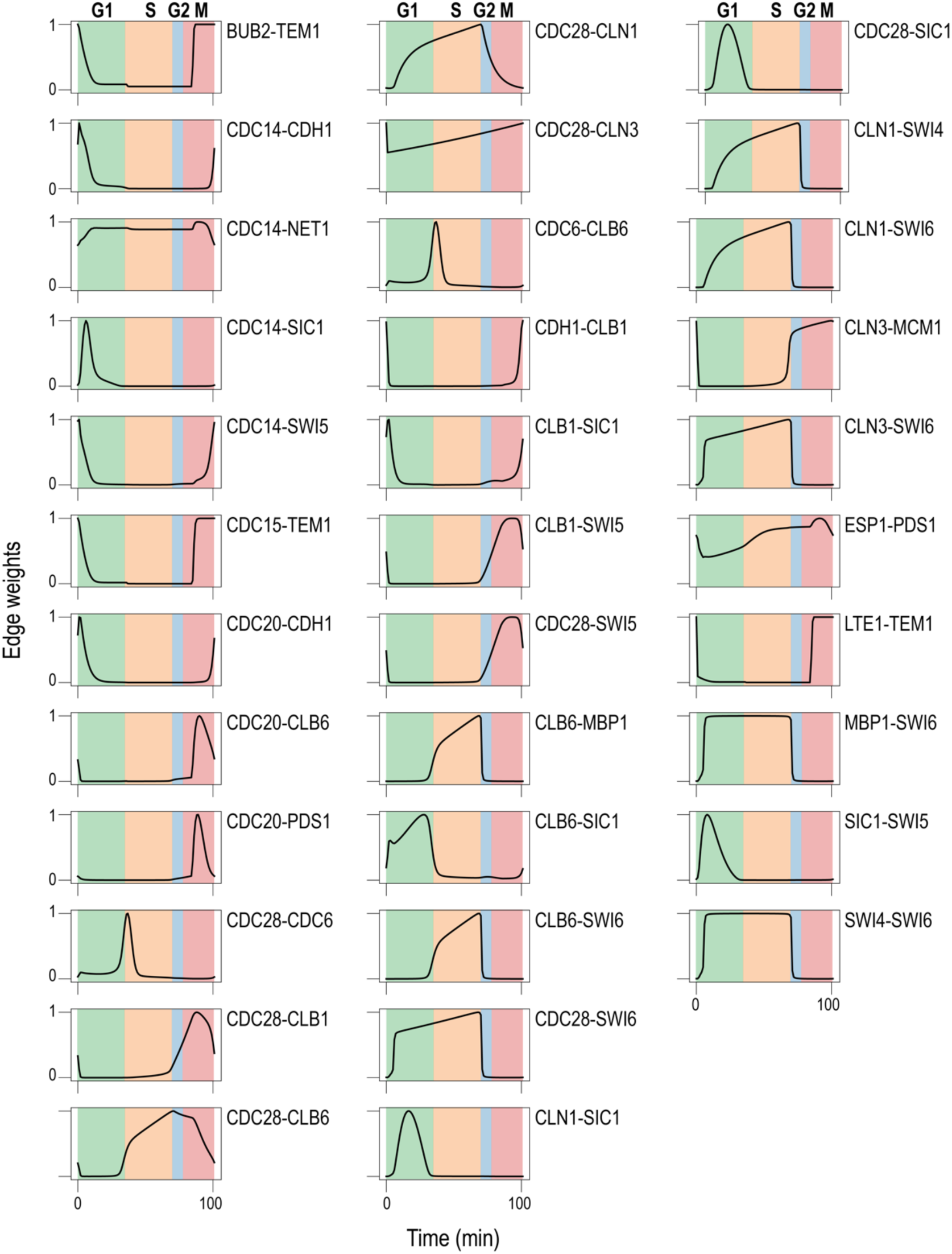
Time series of the weight of all 34 temporal edges in the temporal PPI network (see Figure 2), over one cell cycle. Phases are indicated on top and highlighted with a colour code in each plot.

**Supplementary Figure S2:**
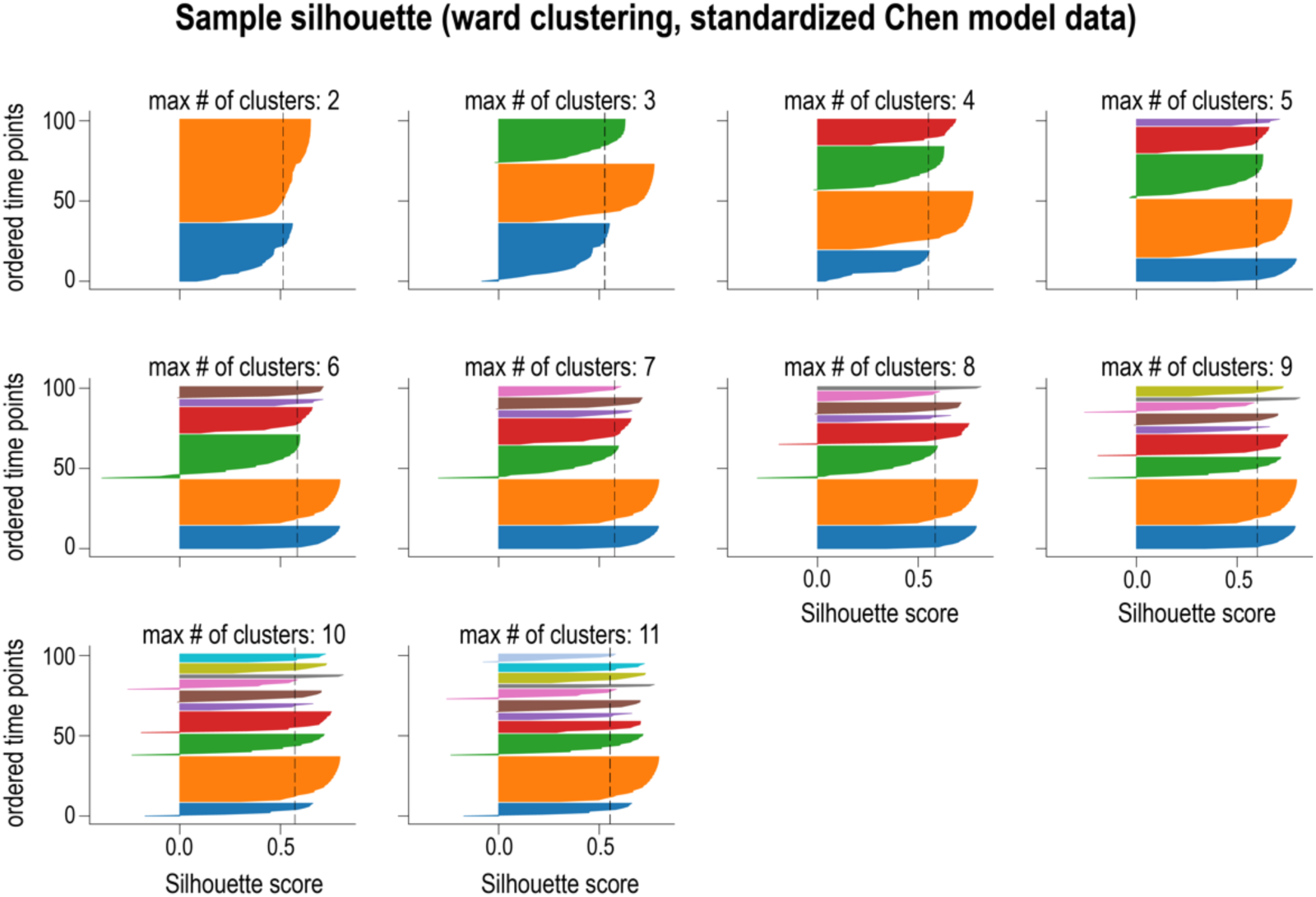
Sample silhouettes, i.e. silhouette score of each snapshot, for the clusters shown in Figure 4 b.

**Supplementary Figure S3:**
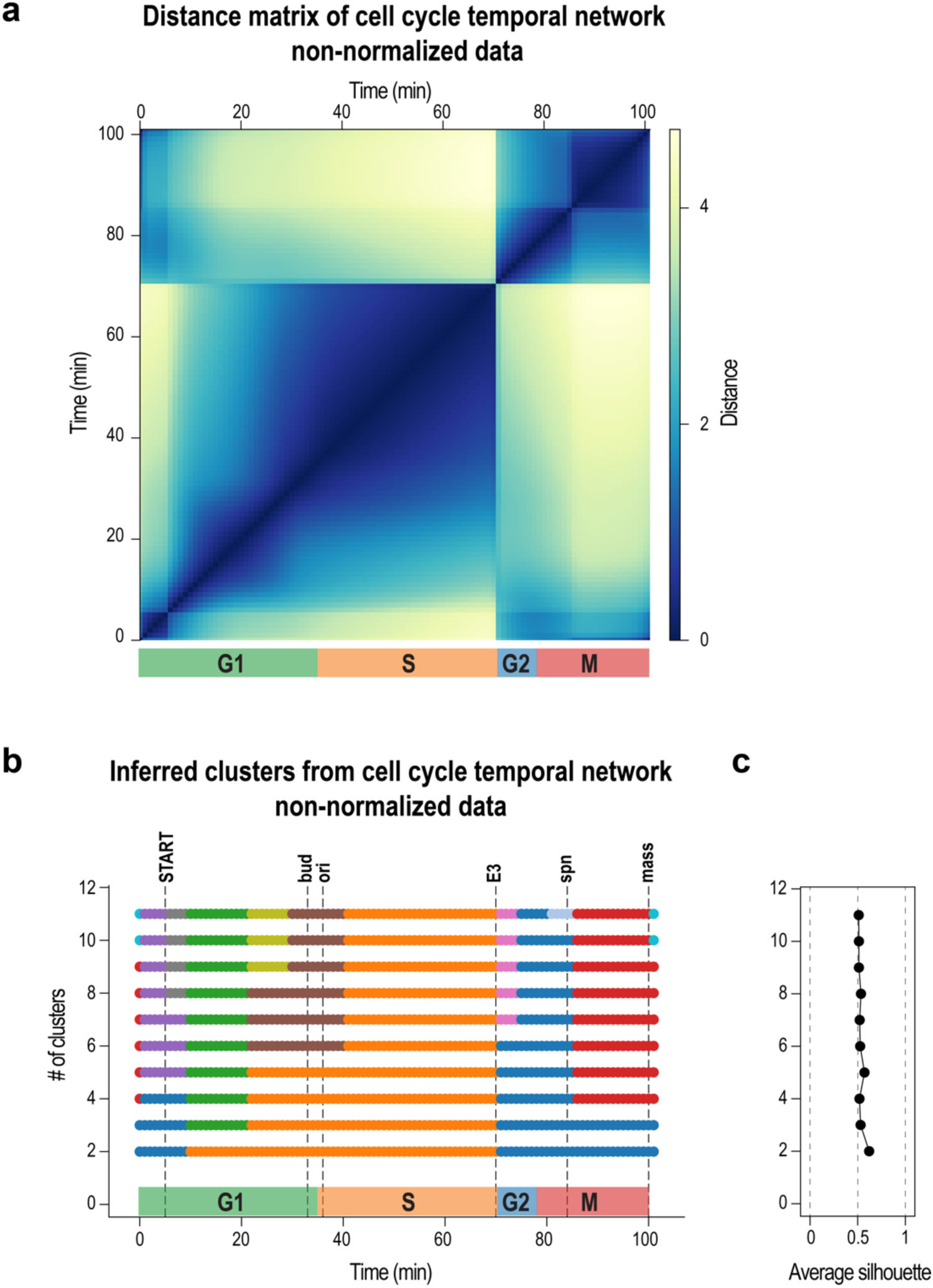
**(a)** Distance matrix **(b)** and inferred clusters with **(c)** their average silhouette score from the cell cycle temporal network using non-normalized edge activities. See also main Figure 4 in the main manuscript.

**Supplementary Figure S4:**
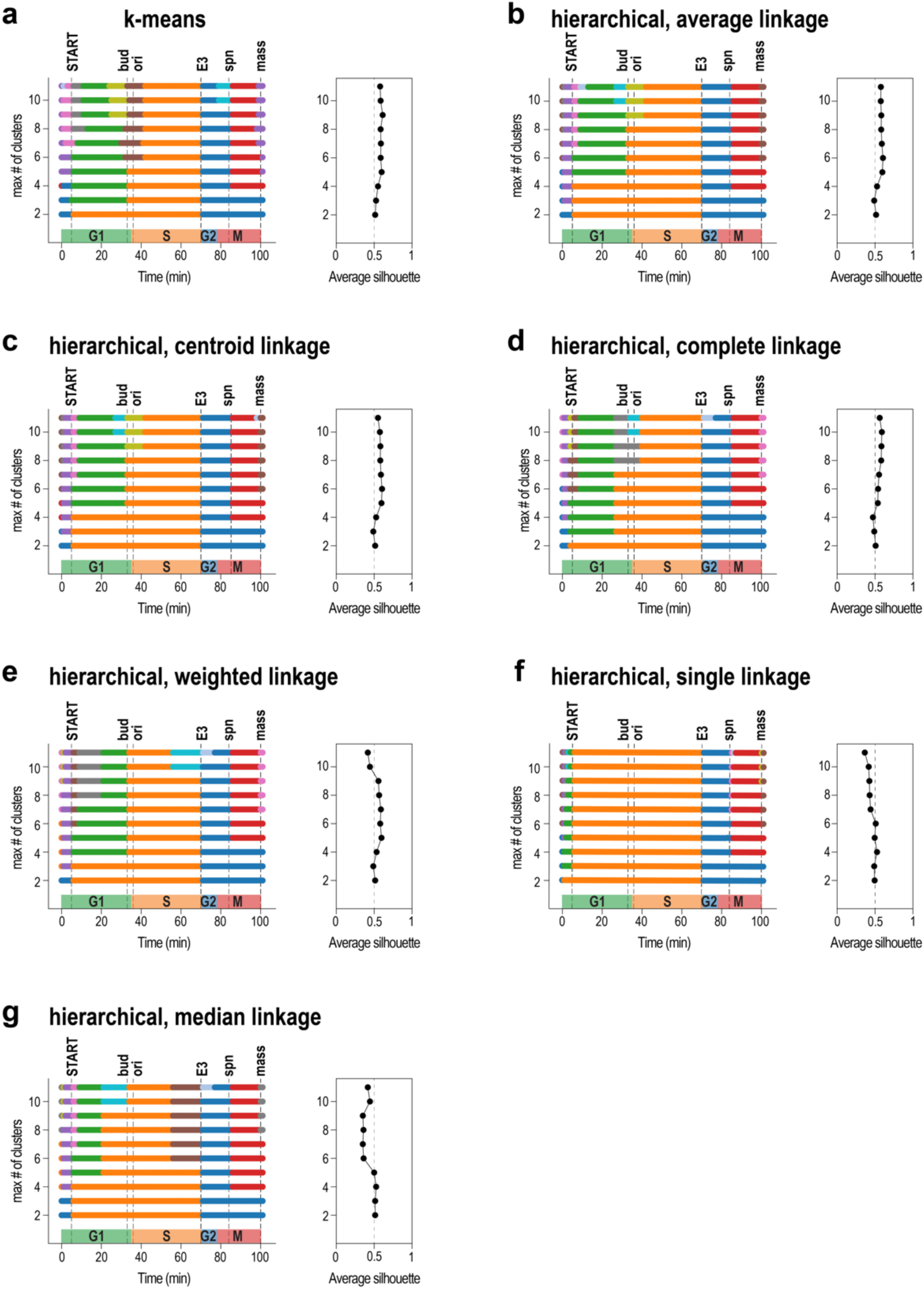
Comparison of our base clustering method, hierarchical clustering with Ward linkage, to 7 other methods: **(a)** k-means, as well as **(b-g)** 6 hierarchical clustering methods. **(b)** Average linkage, **(c)** centroid linkage and **(e)** weighted linkage hierarchical clustering produce similar clusters up to 7 clusters and start to deviate slightly at higher numbers of clusters. **(a)** K-means clustering determines more and earlier deviation at the “bud” and “ori” checkpoints. **(d)** Complete linkage hierarchical clustering tends to cluster a prolonged S phase, with deviation in higher cluster numbers at the “bud” and “ori” checkpoints. **(f)** Single linkage hierarchical clustering fails to find meaningful clusters at higher cluster numbers. It only detects a cluster corresponding to the G1/S phases followed by two clusters corresponding to the two phases G2 and M. Finally, **(g)** median linkage performs similarly to complete linkage at lower numbers of clusters, but then introduces additional clusters towards the end of the S phase at the E3 excitation point.

**Supplementary Figure S5:**
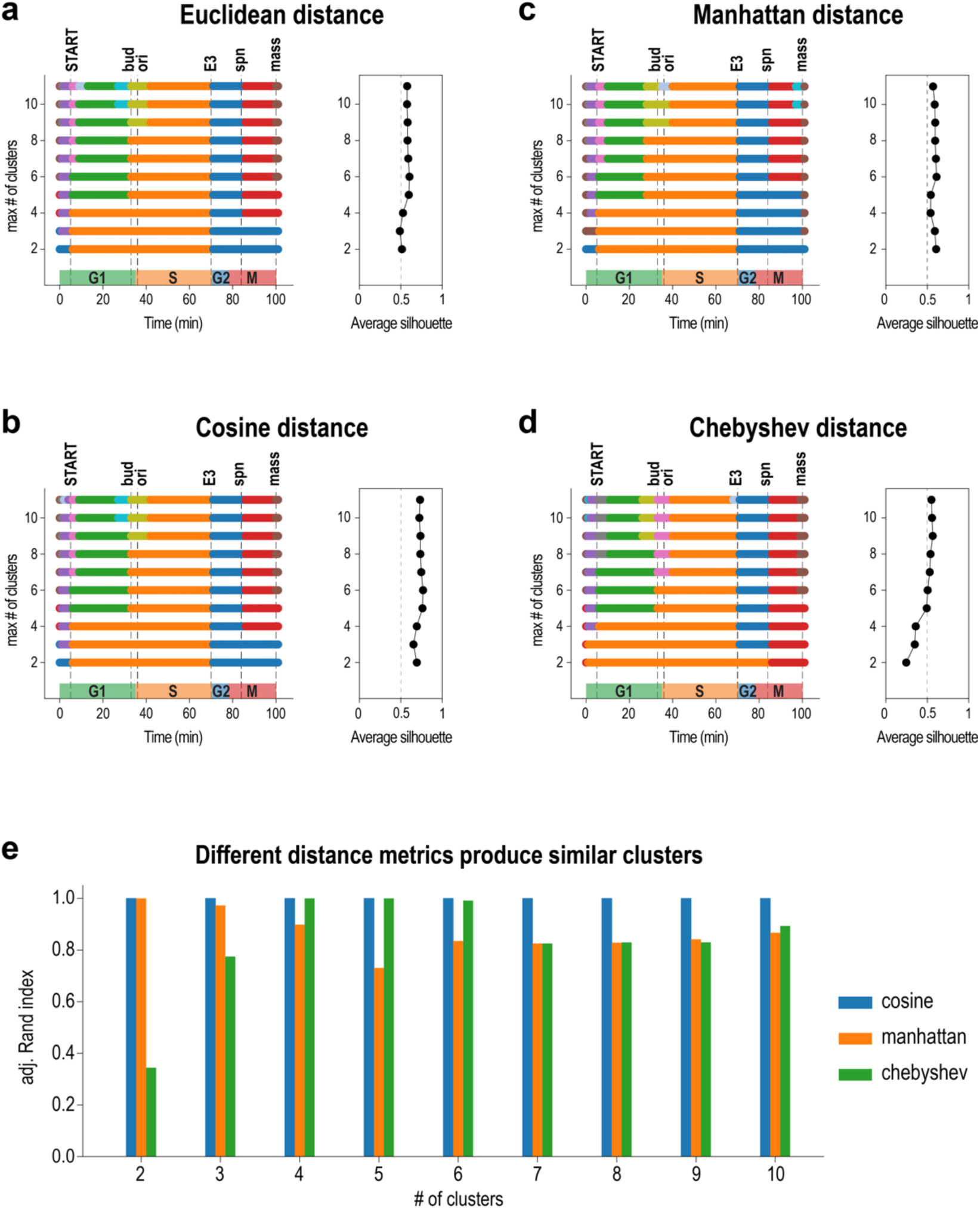
Clustering methods are robust against changes in distance metrics. Comparison of clustering methods obtained with 4 distance metrics: **(a)** Euclidean, **(b)** Cosine, **(c)** Manhattan, and **(d)** Chebyshev. Clustering method: hierarchical with average linkage. Note: Ward linkage requires the Euclidean distance metric to be used. **(e)** Similarity of the 5 clusters obtained by methods in **(b-d)** to those from the Euclidean distance in **(a)**.

**Supplementary Figure S6:**
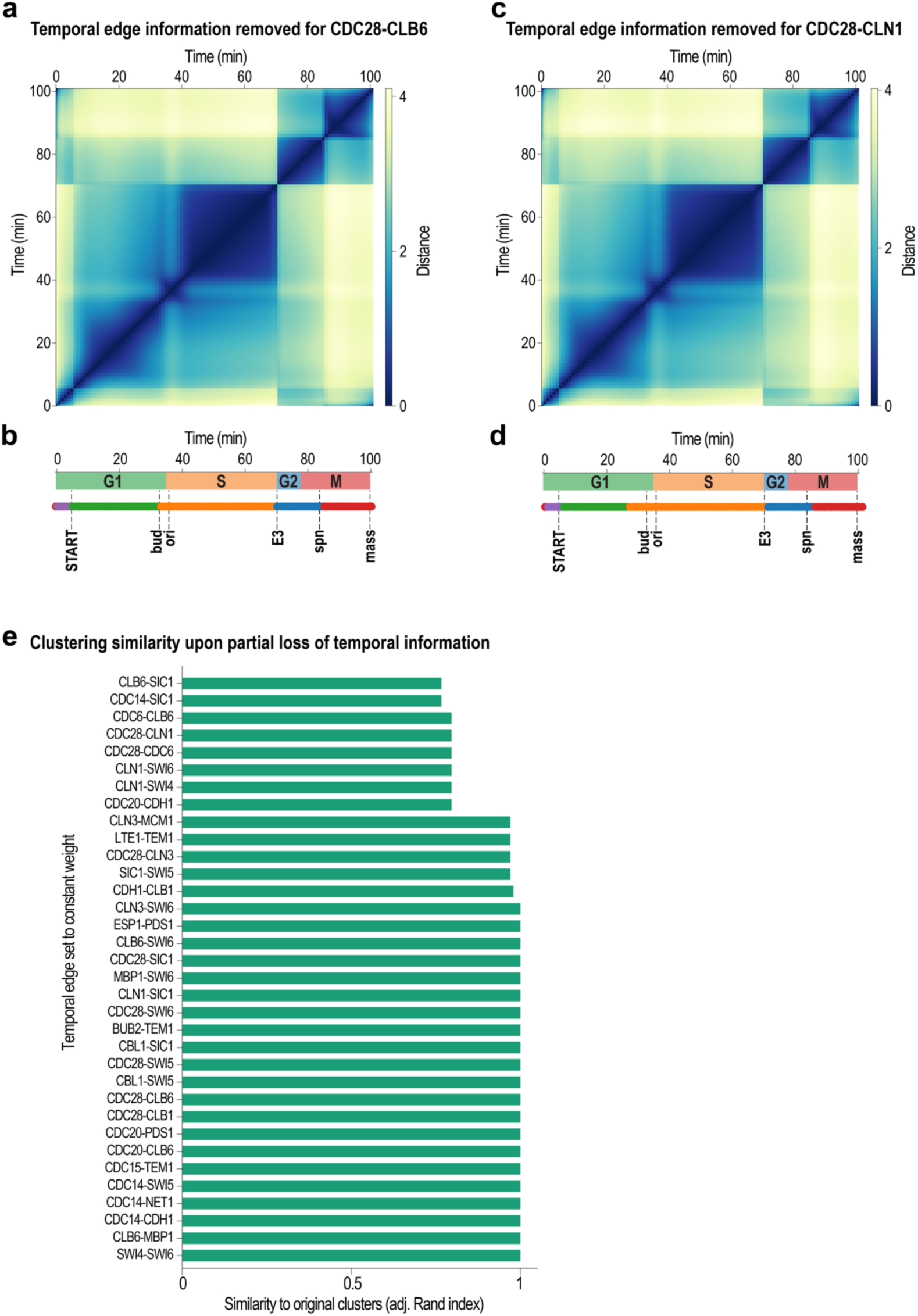
Discarding the temporal information of a random temporal edge does not affect the clustering of the temporal networks into 5 clusters: adjusted Rand Index indicates similarity to the original 5 clusters. Values of 1 indicate identical clusters. The inferred clusters are robust against a randomly chosen edge weight set to a constant value. **(a,b)** Temporal edge information was removed only for CDC28-CLB6 or **(c,d)** CDC28-CLN1. **(e)** Similarity of clustering for single edges set to constant weight.

**Supplementary Figure S7:**
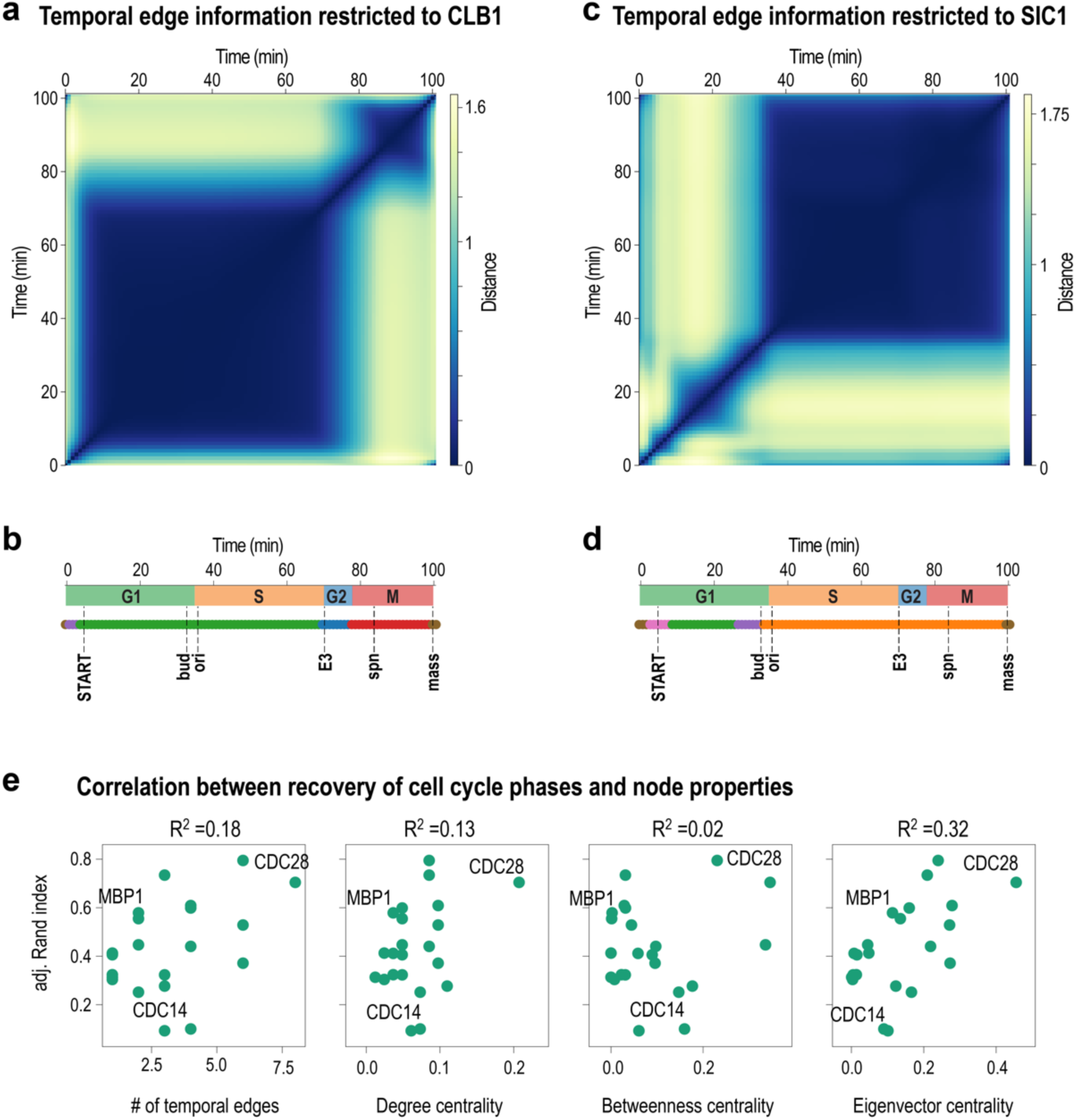
Temporal edge information for some nodes only is not sufficient to infer the original clusters. **(a)** Distance matrix and **(b)** inferred clusters from a temporal network in which temporal information is only kept for the edges of CLB1. One large cluster spanning phases G1 and S was obtained, and the G2/M cluster was detected. **(c)** Distance matrix and **(d)** inferred clusters when temporal information is only kept for the edges of SIC1. The situation was reversed, with a large cluster corresponding to phases S, G2 and M, and smaller clusters corresponding to phase G1. **(e)** We could not find a clear correlation between the # of temporal edges, or typical static network centrality measures (Degree centrality, Betweenness centrality) and only weak correlation between Eigenvector centrality and the adjusted Rand index that indicates good recovery of initial clusters.

**Supplementary Figure S8:**
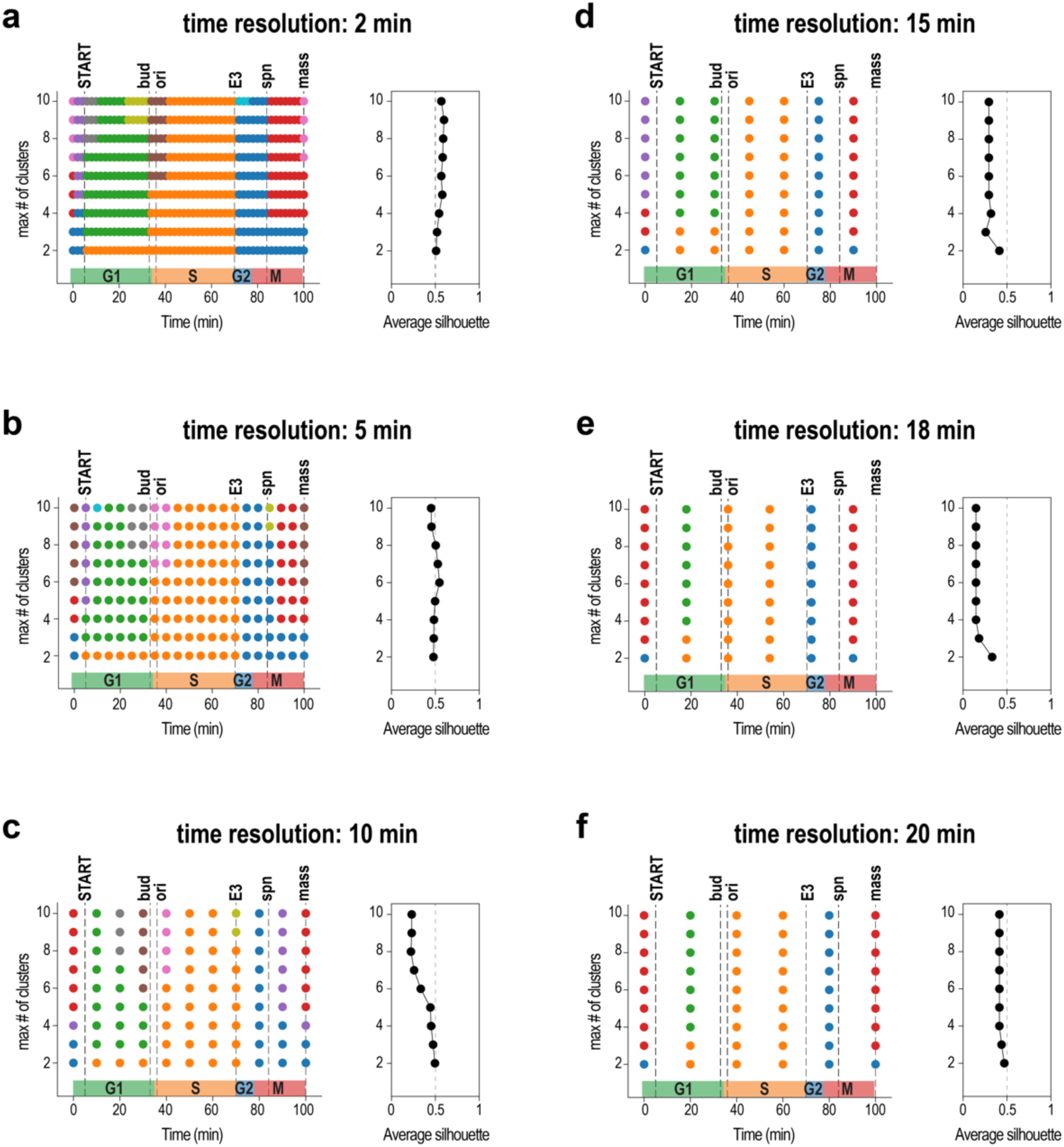
Testing the robustness of the multiscale phase inference using Phasik, with respect to sampling frequency changes. Different sampling frequencies (or equivalently its inverse, the time resolution) of the ODE model provided by Chen were used to build the temporal cell cycle model. **(a)** 2 minutes, **(b)** 5 minutes, **(c)** 10 minutes, **(d)** 15 minutes, **(e)** 18 minutes and **(f)** 20 minutes. The initial 5 phases were well retrieved down to very low sampling frequencies, at which only the maximum obtainable number of clusters was lower.

## References

Alon U. Biological networks: the tinkerer as an engineer. Science. 2003;301(5641):1866–7.

Amoussouvi A, Teufel L, Reis M, Seeger M, Schlichting JK, Schreiber G, et al. Transcriptional timing and noise of yeast cell cycle regulators—a single cell and single molecule approach. Npj Syst Biol Appl. 2018 May 21;4(1):1–10.

Bader JS, Chaudhuri A, Rothberg JM, Chant J. Gaining confidence in high-throughput protein interaction networks. Nat Biotechnol. 2004 Jan;22(1):78–85.

Chechik G, Oh E, Rando O, Weissman J, Regev A, Koller D. Activity motifs reveal principles of timing in transcriptional control of the yeast metabolic network. Nat Biotechnol. 2008;26(11):1251.

Chen KC, Calzone L, Csikasz-Nagy A, Cross FR, Novak B, Tyson JJ. Integrative analysis of cell cycle control in budding yeast. Mol Biol Cell. 2004 Aug;15(8):3841–62.

Chen T, He HL, Church GM. Modeling gene expression with differential equations. In: Biocomputing ’99. WORLD SCIENTIFIC; 1998. p. 29–40. Available from: https://www.worldscientific.com/doi/abs/10.1142/9789814447300_0004

Gelardi V, Godard J, Paleressompoulle D, Claidiere N, Barrat A. Measuring social networks in primates: wearable sensors versus direct observations. Proc R Soc Math Phys Eng Sci. 2020 Apr 29;476(2236):20190737.

Holme P. Modern temporal network theory: a colloquium. Eur Phys J Bs. 2015;88(9):234.

Holme P, Saramäki J. Temporal networks. Phys Rep. 2012;519(3):97–125.

Howard A, Pelc SR. Synthesis of Deoxyribonucleic Acid in Normal and Irradiated Cells and Its Relation to Chromosome Breakage. Heredity. 1953;6:261–73.

Jeong H, Tombor B, Albert R, Oltvai ZN, Barabási A-L. The large-scale organization of metabolic networks. Nature. 2000 Oct;407(6804):651–4.

Kanehisa M, Goto S. KEGG: Kyoto Encyclopedia of Genes and Genomes. Nucleic Acids Res. 2000 Jan 1;28(1):27–30.

Kelliher CM, Leman AR, Sierra CS, Haase SB. Investigating Conservation of the Cell-Cycle-Regulated Transcriptional Program in the Fungal Pathogen, Cryptococcus neoformans. PLOS Genet. 2016 Dec 5;12(12):e1006453.

Koch C, Nasmyth K. Cell cycle regulated transcription in yeast. Curr Opin Cell Biol. 1994 Jun;6(3):451–9.

Komurov K, White M. Revealing static and dynamic modular architecture of the eukaryotic protein interaction network. Mol Syst Biol. 2007;3(1).

Li M, Wu X, Wang J, Pan Y. Towards the identification of protein complexes and functional modules by integrating PPI network and gene expression data. BMC Bioinformatics. 2012;13(1):109.

Li M, Yang J, Wu F-X, Pan Y, Wang J. DyNetViewer: a Cytoscape app for dynamic network construction, analysis and visualization. Bioinforma Oxf Engl. 2018 May 1;34(9):1597–9.

Lopes MA, Zhang J, Krzemiński D, Hamandi K, Chen Q, Livi L, et al. Recurrence quantification analysis of dynamic brain networks. Eur J Neurosci. 2021;53(4):1040–59.

Lord M, Yang MC, Mischke M, Chant J. Cell Cycle Programs of Gene Expression Control Morphogenetic Protein Localization. J Cell Biol. 2000 Dec 25;151(7):1501–12.

Lovrics A, Csikász-Nagy A, Zsély IG, Zádor J, Turányi T, Novák B. Time scale and dimension analysis of a budding yeast cell cycle model. BMC Bioinformatics. 2006 Nov 9;7(1):494.

Masuda N, Holme P. Detecting sequences of system states in temporal networks. Sci Rep. 2019 Dec;9(1):795.

Miritello G, Moro E, Lara R. Dynamical strength of social ties in information spreading. Phys Rev E. 2011 Apr 27;83(4):045102.

Müllner D. Modern hierarchical, agglomerative clustering algorithms. ArXiv11092378 Cs Stat. 2011 Sep 12; Available from: http://arxiv.org/abs/1109.2378

Murray AW. Recycling the Cell Cycle: Cyclins Revisited. Cell. 2004 Jan 23;116(2):221–34.

Nasmyth K. Control of the yeast cell cycle by the Cdc28 protein kinase. Curr Opin Cell Biol. 1993 Apr 1;5(2):166–79.

Nasmyth K. At the heart of the budding yeast cell cycle. Trends Genet. 1996 Oct 1;12(10):405–12.

Newman M. Networks. Second Edition. Oxford, New York: Oxford University Press; 2018.

Ou-Yang L, Dai D-Q, Li X-L, Wu M, Zhang X-F, Yang P. Detecting temporal protein complexes from dynamic protein-protein interaction networks. BMC Bioinformatics. 2014;15(1):335.

Pavlopoulos GA, Secrier M, Moschopoulos CN, Soldatos TG, Kossida S, Aerts J, et al. Using graph theory to analyze biological networks. BioData Min. 2011 Apr 28;4(1):10.

Pedregosa F, Varoquaux G, Gramfort A, Michel V, Thirion B, Grisel O, et al. Scikit-learn: Machine Learning in Python. J Mach Learn Res. 2011;12(85):2825–30.

Pedreschi N, Bernard C, Clawson W, Quilichini P, Barrat A, Battaglia D. Dynamic core-periphery structure of information sharing networks in entorhinal cortex and hippocampus. Netw Neurosci. 2020 Jan 1;4(3):946–75.

Pereira-Leal JB, Enright AJ, Ouzounis CA. Detection of functional modules from protein interaction networks. Proteins Struct Funct Bioinforma. 2004;54(1):49–57.

Pierrelée M, Reynders A, Lopez F, Moqrich A, Tichit L, Habermann B. TimeNexus: A Novel Cytoscape App to Analyze Time-Series Data Using Temporal MultiLayer Networks (tMLNs). 2020 Dec 31; Available from: https://www.researchsquare.com

Przytycka TM, Singh M, Slonim DK. Toward the dynamic interactome: it’s about time. Brief Bioinform. 2010;11(1):15–29.

Rousseeuw PJ. Silhouettes: A graphical aid to the interpretation and validation of cluster analysis. J Comput Appl Math. 1987 Nov 1;20:53–65.

Saramäki J, Moro E. From seconds to months: an overview of multi-scale dynamics of mobile telephone calls. Eur Phys J B. 2015 Jun 24;88(6):164.

Traynard P, Fauré A, Fages F, Thieffry D. Logical model specification aided by model-checking techniques: application to the mammalian cell cycle regulation. Bioinforma Oxf Engl. 2016 Sep 1;32(17):i772–80.

Virtanen P, Gommers R, Oliphant TE, Haberland M, Reddy T, Cournapeau D, et al. SciPy 1.0: fundamental algorithms for scientific computing in Python. Nat Methods. 2020 Mar;17(3):261–72.

Vodermaier HC. APC/C and SCF: Controlling Each Other and the Cell Cycle. Curr Biol. 2004 Sep 21;14(18):R787–96.

Wallach T, Schellenberg K, Maier B, Kalathur RKR, Porras P, Wanker EE, et al. Dynamic Circadian Protein–Protein Interaction Networks Predict Temporal Organization of Cellular Functions. PLOS Genet. 2013 Mar 28;9(3):e1003398.

